# The structural influence of the oncogenic driver mutation N642H in the STAT5B SH2 domain

**DOI:** 10.1101/2024.09.09.612134

**Authors:** Liam Haas-Neill, Deniz Meneksedag-Erol, Ayesha Chaudhry, Masha Novoselova, Qirat F. Ashraf, Elvin D. de Araujo, Derek J. Wilson, Sarah Rauscher

## Abstract

The point mutation N642H of the signal transducer and activator of transcription 5B (STAT5B) protein is associated with aggressive and drug-resistant forms of leukemia. This mutation is thought to promote cancer due to hyperactivation of STAT5B caused by increased stability of the active, parallel dimer state. However, the molecular mechanism leading to this stabilization is not well understood as there is currently no structure of the parallel dimer. To investigate the mutation’s mechanism of action, we conducted extensive all-atom molecular dynamics simulations of multiple oligomeric forms of both STAT5B and STAT5B^*N*642*H*^, including a model for the parallel dimer. The N642H mutation directly affects the hydrogen bonding network within the phosphotyrosine (pY)-binding pocket of the parallel dimer, enhancing the pY-binding interaction. The simulations indicate that apo STAT5B is highly flexible, exploring a diverse conformational space. In contrast, apo STAT5B^*N*642*H*^ accesses two distinct conformational states, one of which resembles the conformation of the parallel dimer. The simulation predictions of the effects of the mutation on structure and dynamics are supported by the results of hydrogen-deuterium exchange (HDX) mass spectrometry measurements carried out on STAT5B and STAT5B^*N*642*H*^ in which a phosphopeptide was used to mimic the effects of parallel dimerization on the SH2 domain. The molecular-level information uncovered in this work contributes to our understanding of STAT5B hyperactivation by the N642H mutation and could help pave the way for novel therapeutic strategies targeting this mutation.

## Introduction

The signal transducer and activator of transcription (STAT) family of proteins are important therapeutic targets for cancer, as well as other diseases ^1–4^. The STAT family comprises seven members: STAT1, STAT2, STAT3, STAT4, STAT5A, STAT5B, and STAT6 ^1;5^. STAT5A and STAT5B are two isoforms of STAT5, which have over 90% sequence identity ^6;7^. In healthy cells, STAT proteins play critical roles in various cellular processes, including cell growth, differentiation, apoptosis, and proliferation ^3;8^. Abnormal regulation of STAT proteins can lead to dysregulation of these cellular processes, which in turn can lead to cancer ^8^. Dysregulation of all STAT proteins is reported in various types of human cancer, and hyperactivation of STAT3 and STAT5 (including both the STAT5A and STAT5B isoforms) is frequently observed in cancer cells ^1;2;5;8^.

STAT proteins are activated in response to extracellular signals via the JAK/STAT pathway ^3–5;11;12^. Prior to phosphorylation by Janus kinases (JAKs), STAT proteins exist in an equilibrium between monomer and antiparallel dimer forms (Figure 1a,c) ^8;9;13^. The conventional mode of STAT activation occurs when extracellular signals activate JAKs, which in turn activate STATs by phosphorylating a conserved tyrosine residue ^3;14^. Phosphorylated STAT proteins then form a parallel dimer (Figure 1e) and enter the cell nucleus, where they bind to DNA and activate transcription of their target genes ^3;12;14^. Although this process is established as the primary, canonical mode of STAT function, other “non-canonical” processes have been observed ^12^. Specifically, STAT proteins have been observed to enter the nucleus in the unphosphorylated state, and STATs 1, 2, and 3 can activate transcription without being phosphorylated ^12^. Unphosphorylated STAT5A (as well as STAT5B) can function as a transcriptional repressor ^12;15;16^. In solution, unphosphorylated STAT5A is in equilibrium between monomer and antiparallel dimer configurations, with the monomer being the dominant species ^13^. These findings highlight the potential functional roles of multiple oligomerization states of STAT proteins while emphasizing the importance of the parallel dimer state.

**Figure 1:**
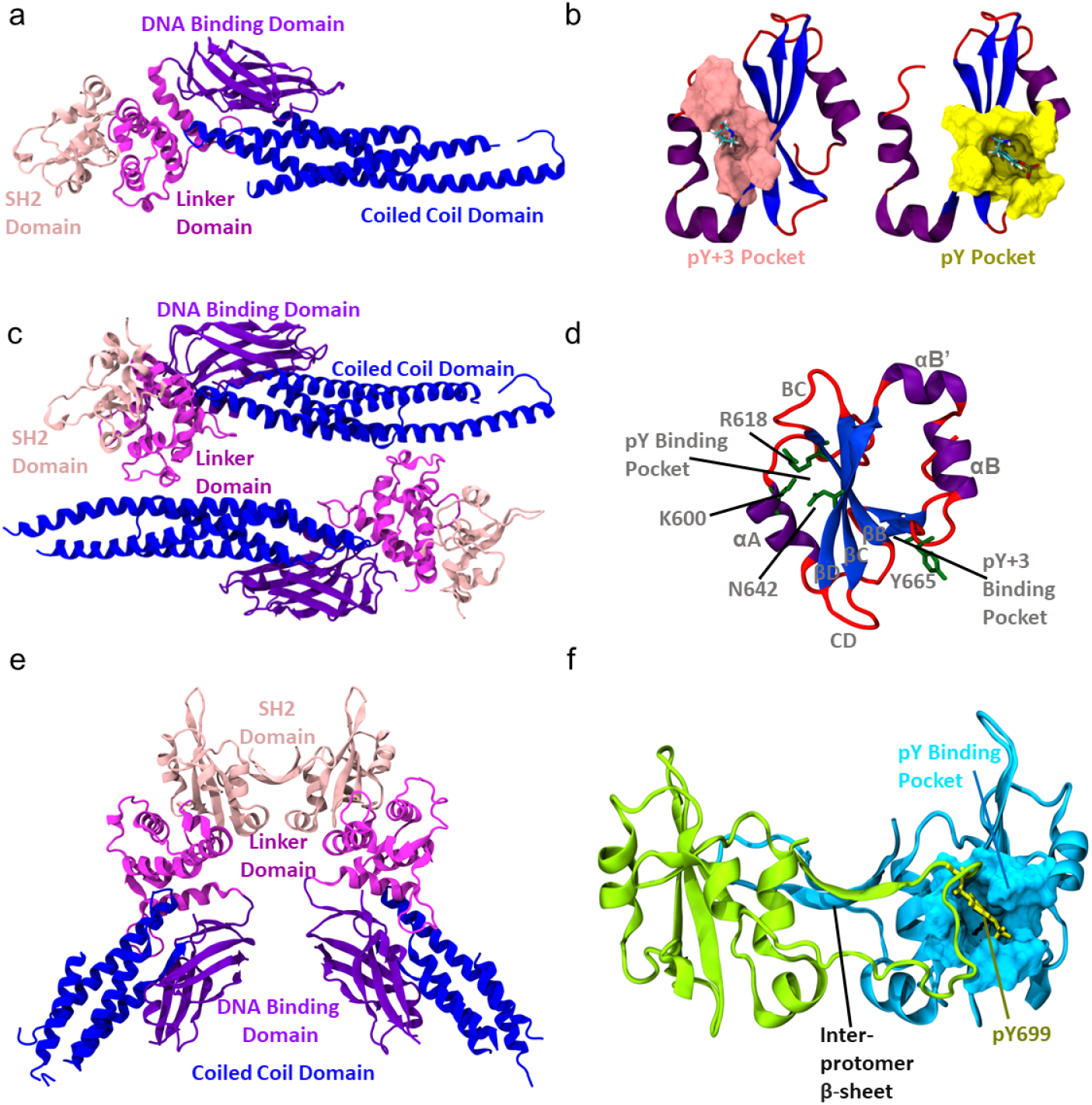
3D structure of STAT5B and its SH2 domain. (a) Structure of the STAT5B monomer core fragment shown in cartoon representation with each domain labelled (coiled-coil domain: 138-331, DNA binding domain: 332-471, linker domain: 498-591, SH2 domain 592-685). The structure shown is chain B from the crystal structure of the STAT5B antiparallel dimer, PDB 6MBZ ^9^. (b) 3D structures of the SH2 domain from STAT5B (pY and pY+3 pockets in surface representation; pY and pY+3 residues in licorice representation). (c) Structure of the STAT5B antiparallel dimer core fragment (PDB 6MBZ ^9^) shown in cartoon representation with each domain labelled. (d) SH2 domain with the secondary structure elements labelled. Key residues are shown in licorice representation. (e) Structure of the STAT6 parallel dimer core fragment (PDB 4Y5U ^10^. (f) Structure of the STAT5B parallel dimer SH2 domain and phosphotyrosine motif region (including residues 589-709). Residue pY699 from chain A (green, licorice representation) interacts with the pY binding pocket of chain B (blue, surface representation). The structure of the SH2 domain shown in panels (b), (d) and (f) corresponds to the SH2 domain in the homology model of the parallel dimer. The simulations of the homology model also include the linker domain, which is not shown here for clarity. The crystal structures of STAT5B and STAT6 in (a), (c), and (e) include only the core fragment and are missing the N-terminal oligomerization and transactivation domains.

While the STAT proteins carry out diverse functions ^3;12^, they share a similar structure that comprises six domains (Figure S1a) ^3;8;9;11^. Four ordered domains constitute the core fragment (Figure 1a): the coiled-coil domain, DNA binding domain, linker domain, and the SH2 domain ^9;10;17^. There are two additional domains: the N-terminal domain and the transactivation domain ^9^. The N-terminal domain (or oligomerization domain) mediates antiparallel dimer formation ^18^, as well as the formation of tetramers ^11;19^. The transactivation domain is predicted to be disordered in all human STAT proteins (refer to Figure S2 for the disorder prediction obtained using Metapredict V2 ^20;21^). The SH2 domain and the transactivation domain are separated by a short sequence of residues containing the phosphorylated tyrosine (pY) residue that we refer to as the phosphotyrosine motif region. The SH2 domain (Figure 1d) facilitates interactions that stabilize the parallel dimer form of the protein ^9;10^. The pY residue in the phosphotyrosine motif region of each protomer interacts with the phosphopeptide binding pocket (pY pocket, Figure 1b) of the conjugate protomer ^9^. This mutual interaction forms the basis of the parallel dimer interface (Figure 1b,f). Mutual interaction between the pY+3 residue and the specificity pocket (pY+3 pocket, Figure 1b) also contributes to the parallel dimer interface and determines affinity for specific binding partners ^22^.

The majority of disease-causing mutations in STAT proteins occur in the SH2 domain ^4^. Multiple oncogenic driver mutations occur in the SH2 domain of STAT5B, with N642H being the most frequently observed ^9;23–25^. The N642H mutation is linked to aggressive forms of leukemia ^26^, increased risk of relapse ^27^, and drug resistance ^9;28^. This mutation has also been reported to stabilize STAT5B parallel dimers, prolong or increase phosphorylation ^22;23;29^, increase parallel dimer binding affinity ^23^, and render STAT5B constitutively active ^27;30^. The increased propensity to form parallel dimers results in hyperactivation ^23;27;30^. However, the molecular underpinnings of constitutive and prolonged phosphorylation are not well understood because no structure is available for the STAT5B parallel dimer.

Molecular dynamics (MD) simulations can be used to study the effect of mutations on protein structure and dynamics, which is especially useful in cases where experimental data is challenging to obtain ^31–34^. Molecular simulations have been used to study the structure and dynamics of SH2 domains ^35–38^ and STAT proteins ^9;39–42^. Several of these studies have focused on STAT5, including the STAT5B monomer and antiparallel dimer, as well as the STAT5A parallel dimer ^9;39;41;43^. Computational methods have also been used to model the structure and dynamics of STAT5A bound to a phosphopeptide ^42^. However, there is currently no crystal structure or model of the STAT5B parallel dimer or molecular-level information on the dynamics of this system. A structural and dynamic picture of the STAT5B parallel dimer interface and how the N642H mutation alters it could accelerate the rational design of novel therapeutics that target this mutation.

Here, we carry out MD simulations with a cumulative sampling time of nearly 100 *µ*s to reveal structural and dynamic differences between the STAT5B and STAT5B^*N*642*H*^ proteins. The N642H mutation changes the hydrogen bonding network within the pY pocket; specific inter-protein hydrogen bonds are more probable in the STAT5B^*N*642*H*^ parallel dimer. These changes could explain the experimentally observed increase in dimer stability and prolonged phosphorylation with the N642H mutation ^23;29^. Furthermore, the results of our simulations of the monomers and antiparallel dimers indicate that the N642H mutation causes a rigidification of the SH2 domain, even in the absence of phosphotyrosine binding. The STAT5B^*N*642*H*^ antiparallel dimer samples two states, one exhibiting a marked similarity in structure and dynamics to the pY-bound parallel dimer. These two states are also observed in MD simulations of the SH2 and linker domains of two other proteins from the STAT family, STAT1 and STAT5A. Finally, we perform hydrogen-deuterium exchange (HDX) measurements that indicate both phosphopeptide binding and the N642H mutation cause a rigidification of the STAT5B SH2 domain compared to apo wild type, consistent with what we observe using simulation. Together, our results provide a structural and dynamic view of the N642H mutation’s influence on the STAT5B SH2 domain and the parallel dimer interface.

## Results

To investigate the influence of the N642H mutation on the structure and dynamics of STAT5B, we performed MD simulations of STAT5B in monomeric form and different dimerization states, both with and without the N642H mutation (refer to Table 1 and Table S1 for a list of simulations). Because no experimental structure is available for the STAT5B parallel dimer, we constructed a homology model (Figure 1f) using the crystal structure of the STAT6 parallel dimer (Figure 1e) ^10^ as a template. STAT6 was chosen because it is the STAT protein with the most similar sequence to STAT5B for which a crystal structure of the parallel dimer is available. We included only the linker and SH2 domains in our simulations of the homology model of the parallel dimer. We simulated this system because there are no inter-protomer interactions involving the DNA binding domain or the coiled coil domain in the crystal structure of the parallel dimer of STAT6 ^10^. A list of all published crystal structures of mammalian STAT proteins in different oligomerization states is provided in Table S2.

**Table 1:** List of simulations performed in this study.

| System | # Trajectories | Sampling Time (μs) | # atoms |
| --- | --- | --- | --- |
| STAT5B parallel dimer<br>linker + SH2 | 3 | 12 | 110k |
| STAT5B antiparallel dimer<br>core fragment | 3 | 6.86 | 415k |
| STAT5B monomer<br>core fragment | 6 | 12 | 375k |
| STAT5B<sup>N642H</sup> parallel dimer<br>linker + SH2 | 3 | 12 | 155k |
| STAT5B<sup>N642H</sup> antiparallel dimer<br>core fragment | 5 | 16.9 | 415k |
| STAT5B<sup>N642H</sup> monomer<br>core fragment | 6 | 12 | 375k |
| STAT1 monomer<br>linker + SH2 | 6 | 15.772 | 70k |
| STAT5A monomer<br>linker + SH2 | 6 | 12 | 80k |

### A model for the parallel dimer of STAT5B

Using the set of STAT crystal structures, we constructed a conformational landscape of STAT SH2 domains (see Methods, Figure 2a, Figure S3). The resulting conformational space shows that SH2 domains of different members of the STAT family separate into distinct clusters (Figure 2a). We projected the trajectory of the equilibration simulations starting from the homology model of the parallel dimer onto this space (Figure 2a). Structures from this trajectory are initially found near the STAT6 region, consistent with the fact that a STAT6 structure ^10^ was used as a template for the homology model. As the simulation progresses, the trajectory drifts from its starting location near the STAT6 region towards the STAT5 region. This drift occurs mainly along the second principal component, which describes the proximity of the *β*D-strand to the *β*B-strand and the N-terminus of the protein (see Methods, Figure S4). The similarity between the structure of the equilibrated homology model and the crystal structures of STAT5A and STAT5B supports the utility of this model in describing the parallel dimer state of STAT5B.

**Figure 2:**
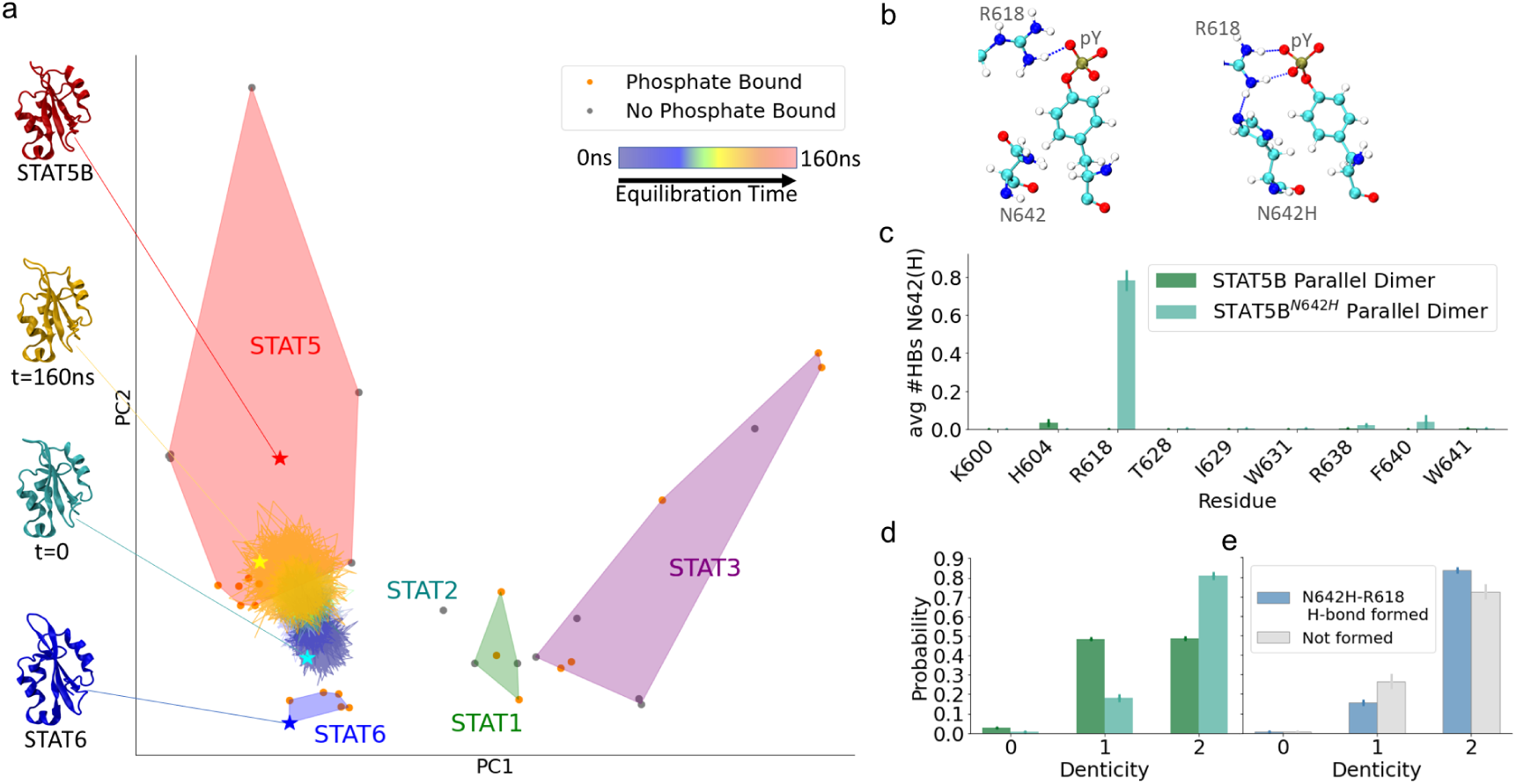
A model for the STAT5B parallel dimer and effects of the N642H mutation on the parallel dimer interface. (a) The conformational landscape of mammalian STAT SH2 domains is shown. The simulation trajectory of the parallel dimer equilibration simulation is projected onto the space, with each 20 ns of the equilibration simulation indicated as a different colour (see legend). Regions of the conformational space encapsulating each STAT protein are indicated as shaded regions. Phosphate-bound crystal structures (including parallel dimers and phosphopeptides) are shown in orange, while apo crystal structures are shown in grey. Key structures from the conformational space are shown along the side, and are indicated on the conformational space by stars with colours matching the structures. The same conformational space with all STAT crystal structures labelled is provided in Figure S3. (b) Representative configuration of residue R618, the pY residue and residue 642 (N642, left; N642H, right) to illustrate the main difference in the hydrogen bonding network in the pY pocket due to the mutation. (c) Average number of hydrogen bonds between the mutation site residue (642) and other SH2 domain residues in simulations of STAT5B (dark green) and STAT5B^*N*642*H*^ (light green) parallel dimers. (d) Probability distribution of the denticity of the pY-R618 hydrogen bond in simulations of STAT5B (dark green) and STAT5B^*N*642*H*^ (light green) parallel dimers. (e) Conditional probability distribution of denticity of the pY-R618 hydrogen bond in the STAT5B^*N*642*H*^ simulations, conditioned on the N642H-R618 hydrogen bond being formed (blue) or not formed (grey). Error bars in panels (c-e) indicate standard error of the mean obtained by treating each SH2 domain in the simulations as an independent measurement.

Over the course of the simulations of the parallel dimer, the pY residue forms contacts with multiple residues (Figure S5). Using the probabilities of these contacts, we determined which residues constitute the pY binding pocket (see Methods). Contacts with residues in the pY pocket persist for nearly 100% of all simulation trajectories of the STAT5B parallel dimer. The fact that we do not see any rearrangement or dissociation of these contacts during the simulation suggests that the interaction between pY and the SH2 domain is stable, which adds to our confidence in the homology model.

In SH2 domains, binding specificity is primarily determined by interactions with residues following the pY residue in the sequence ^44;45^. Without an experimental structure for the STAT5B parallel dimer, the interactions stabilizing the dimeric state and responsible for the binding specificity of this SH2 domain are not known. To determine the primary constituents of the dimer interface, we computed the inter-protomer contact probabilities from the wild-type parallel dimer simulations (Figure S6a,c). Stable interactions are present between the SH2 domain and the pY residue, hydrophobic residues pY+1 (V700) and pY+3 (P702) and less prominent interactions with pY+2 (K701). Another hydrophobic residue in the vicinity, pY+5 (I704), also forms transient contacts with the SH2 domain, but its most prominent contacts stabilize an inter-protomer *β*-sheet formed by residues 703-707 (pY+4 to pY+8, Figure 1f) of both protomers. This inter-protomer *β*-sheet represents a significant portion of the parallel dimer interface. The interactions we observe with residues following the pY residue show that the stability of the parallel dimer interface of STAT5B arises from multiple interactions in addition to those involving the pY and pY+3 binding pockets. This dimer interface is similar to those seen in other STAT proteins. The parallel dimer structures of STAT1^46;47^, STAT3 ^48^, and STAT6 ^10^ also have inter-protomer contacts between antiparallel strands formed by the residues following pY. Hydrophobic interactions between residues pY+1, pY+3, or pY+5 with pockets in the SH2 domain are also typical of SH2 domain-phosphopeptide interactions ^36;44^, consistent with our observations.

### The N642H mutation alters the pY pocket interaction network and hydrogen bonding with pY

To investigate potential causes of the prolonged phosphorylation and increased dimer affinity of STAT5B^*N*642*H*^, we focused on changes in interactions at the pY-SH2 binding interface. Specifically, we characterized van der Waals contacts (Figure S6a,b, Figure S7b) and the hydrogen bonding network (Figure 2b-e, Figure S7a) in the pY pocket. The N642H mutation changes the hydrogen bonds both within the pY-binding pocket (Figure 2b,c) and between the pocket and the pY residue (Figure 2d, Figure S7a) in the STAT5B parallel dimer. The histidine residue at position 642 forms hydrogen bonds with the highly conserved ^49^ residue R618 (Figure 2b,c, Figure 1d). Hydrogen bonds between residues N642 and R618 are virtually absent in the wild-type dimer (Figure 2c). When a hydrogen bond is formed between H642 and R618, R618 is in a more optimal orientation to form contacts with the pY residue. As a result, hydrogen bond formation between pY and R618 increases on average (Figure S7a). Specifically, we see a significant increase in the probability of bidentate hydrogen bonding between pY and R618 in STAT5B^*N*642*H*^ (Figure 2d), particularly when the N642H-R618 hydrogen bond is formed (Figure 2e). We also observe a lower prevalence of hydrogen bonds between S620 and S622 with pY in the STAT5B^*N*642*H*^ parallel dimer, although bidentate hydrogen bonds still form between pY and S622.

An arginine residue at this position (corresponding to R618 in STAT5B) is the most important determinant of pY recognition ^50;51^ and forms a strong bidentate hydrogen bond with the pY residue ^52^ in other SH2 domains. Arginine is highly conserved at this position across all SH2 domains, occurring in 118 out of 121 human SH2 domain sequences ^49^. Furthermore, histidine occurs at position 642 in 80 of these sequences ^49^, but never occurs in the SH2 domains that lack arginine in position 618 ^53^. This conditional conservation of H642 on R618 hints at the importance of their mutual interaction to strengthen pY coordination. It also supports the hypothesis that the N642H mutation strengthens pY binding via reorienting R618 to form strong hydrogen bonds with pY. Structures of other SH2 domains ^54;55^ containing arginine and histidine at these sites also have hydrogen bonds between these residues, as well as hydrogen bonds between R618 and pY. Additionally, the importance of hydrogen bonding between H642 and R618 has been reported in other SH2 domains for optimally positioning the guanidinium group of R618 for pY binding ^55;56^. Together, these observations of the sequence conservation and structure of other SH2 domains in prior studies are consistent with the differences in the hydrogen bonding network we observe between parallel dimers due to the N642H mutation.

We next investigated the influence of the N642H mutation on contacts at the dimer interface. Specifically, we computed the number of protein atoms in contact with the pY residue (Figure S7b). The results of this analysis show that the N642H mutation, on average, significantly increases the average number of atoms in contact between the binding pocket residues and pY, particularly involving residues in the *β*D-strand, but also with most other pocket residues. These additional contacts could contribute to the increased parallel dimer affinity of STAT5B^*N*642*H*^. We also observe differences in non-specific contacts between N642(H) and *α*A, specifically with residue H604 (Figure S8a). These additional contacts could serve to stabilize the positioning of N642H and, therefore, contribute indirectly to stabilizing the orientation of R618 in STAT5B^*N*642*H*^.

An additional difference we see in the parallel dimers of STAT5 and STAT5B^*N*642*H*^ is that the phosphotyrosine motif region of the STAT5B^*N*642*H*^ parallel dimer samples a different conformation than that of the wild type (Figure S9a-c). The inter-protomer *β*-sheet is significantly less probable in STAT5B^*N*642*H*^ (Figure S9a), and several interactions between the two protomers are different, including significantly reduced contact between the pY+3 residue and residues 706-709 (Figure S6a,b, S9b,c). We also see increased contact between the pY+5 residue and the SH2 domain (Figure S6a,b). The structural differences due to the mutation in the phosphotyrosine motif region could be due to this region’s increased propensity for disorder compared to the rest of the protein.

To investigate this possibility further, we reviewed what is known about the structure of the phosphotyrosine motif region. First, while it is present in the STAT5B construct used for crystallization, most of the phosphotyrosine motif region (residues 687-703) is not resolved in the only available crystal structure of STAT5B ^9^. Using Metapredict V2 ^20;21^, we obtained a prediction for the propensity of each residue to be disordered in human STAT proteins (Figure S2). The phosphotyrosine motif region is predicted to be disordered in STAT6 and above the baseline in STAT5B (although not above the threshold to be labelled as disordered). This region is predicted to be disordered in STAT1, STAT3 and STAT4 and ordered in STAT2 and STAT5A. Both our simulation results and the crystal structure of STAT6 ^10^ show the phosphotyrosine motif region as largely structured in the parallel dimer. We therefore propose that the phosphotyrosine region may be an intrinsically disordered region that undergoes folding on binding to the other protomer in the parallel dimer, but is disordered in the monomeric state. Disordered or flexible proteins may require more sampling time to adequately explore their conformational space ^57^. Their shallow energy landscapes with many minima also leave them more easily influenced by perturbations. Since the phosphotyrosine motif region is more disordered than the rest of the SH2 domain, it is possible that perturbations, such as the N642H mutation, may destabilize this region more than other regions of the SH2 domain. Therefore, we acknowledge the limitations of the above analysis. Furthermore, the fact that the inter-protomer *β*-sheet dissociates in the simulations with the mutation could result from introducing a mutation into a homology model, which may not fully capture the effects of the mutation on the phosphotyrosine region due to limitations in sampling and the model.

Other explanations for the molecular mechanism of the N642H mutation have been suggested in previous studies. Fahrenkamp et al. suggested that *π*-*π* interactions may be present between the aromatic rings of histidine at position 642 and phosphotyrosine ^41^. We, therefore, analyzed the pY-N642H ring geometries in our simulations (Figure S10). The two rings tend to have a relative stacking angle between 70° −90 ° and a centroid distance larger than 6 *Å*. This distance is larger than the optimal distance for His-Tyr *π*-*π* interactions ^58;59^. While it is possible that *π*-*π* interactions do contribute to the interaction between STAT5B^*N*642*H*^, our data suggest that it does not play a dominant role. We note, however, that *π*-*π* stacking geometries are not well-reproduced by MD simulations ^60^. A molecular docking study suggested that the improved parallel dimerization affinity in STAT5B^*N*642*H*^ stems from improved electrostatic interaction between N642H and pY ^23^. That work does not indicate that N642(H)-R618 interactions play a major role, but this may be due to the limitations of docking models. Our simulations do suggest that N642H has increased atomic contacts with pY (Figure S7b). The additional changes in N642(H)-R618 interactions due to the mutation are likely only accessible because MD simulations allow for larger conformational changes, which would not be captured using docking methods.

Together, the stabilization of the R618-pY hydrogen bonding interaction and the increase in the van der Waals interactions between pY and the binding pocket indicate that STAT5B^*N*642*H*^ binds the pY residue more tightly than the wild-type STAT5B. We propose that these stabilizing interactions in the parallel dimer lead to the prolonged phosphorylation observed for STAT5B^*N*642*H*^. Notably, however, neither the increase in N642(H)-H604 contact nor the increase in N642(H)-R618 contact is observed in the apo simulations (Figure S8a). In fact, we observe that both N642(H)-H604 (Figure S8a) and N642(H)-R618 hydrogen bonds (Figure S11) are slightly less frequent in apo STAT5B^*N*642*H*^ compared to wild type. This could indicate that the structural differences contributing to increased parallel dimerization are only present upon a conformational change induced by phosphopeptide binding. While we observe structural differences involving residues 706-709 between wild-type and N642H ensembles (Figure S6a,b), these differences likely arise because this region is disordered, as discussed above. We do not think these differences contribute to the functional differences between wild-type STAT5B and STAT5B^*N*642*H*^. Furthermore, our results provide evidence that the increased oncogenicity of STAT5B^*N*642*H*^ is not due to simple electrostatic interactions or *π*-*π* stacking between N642H and pY. These predictions provide a plausible structural mechanism of STAT5B hyperactivation by the N642H mutation.

### The N642H mutation rigidifies the SH2 domain

We next investigated if the differences in the pY pocket interactions described above translate to broader structural differences. Specifically, we computed the volume of the pY and pY+3 binding pockets. The distributions of the volume of the pY pocket are similar, but the pY+3 pocket in apo STAT5B^*N*642*H*^ is more compact on average compared to apo STAT5B (Figure 3a,b). Both apo systems have significantly larger binding pockets than the pY-bound parallel dimer systems (Figure 3a-d). These results indicate that the pY and pY+3 pockets contract due to interactions with the phosphotyrosine motif region of the conjugate protomer. The N642H mutation causes the pY+3 pocket to contract even in the absence of pY binding, although to a lesser extent than in the parallel dimer. Apart from changes in average pY+3 pocket size, the volume of the pY+3 binding pocket is more narrowly distributed in apo STAT5B^*N*642*H*^ compared to apo STAT5B. This indicates that the pY+3 binding pocket of STAT5B^*N*642*H*^ is more rigid than in STAT5B. For comparison, the distribution of the volume of the pY and pY+3 pockets is very narrow in the parallel dimer (Figure 3c,d), indicating that parallel dimerization also leads to rigidification of the binding pockets.

**Figure 3:**
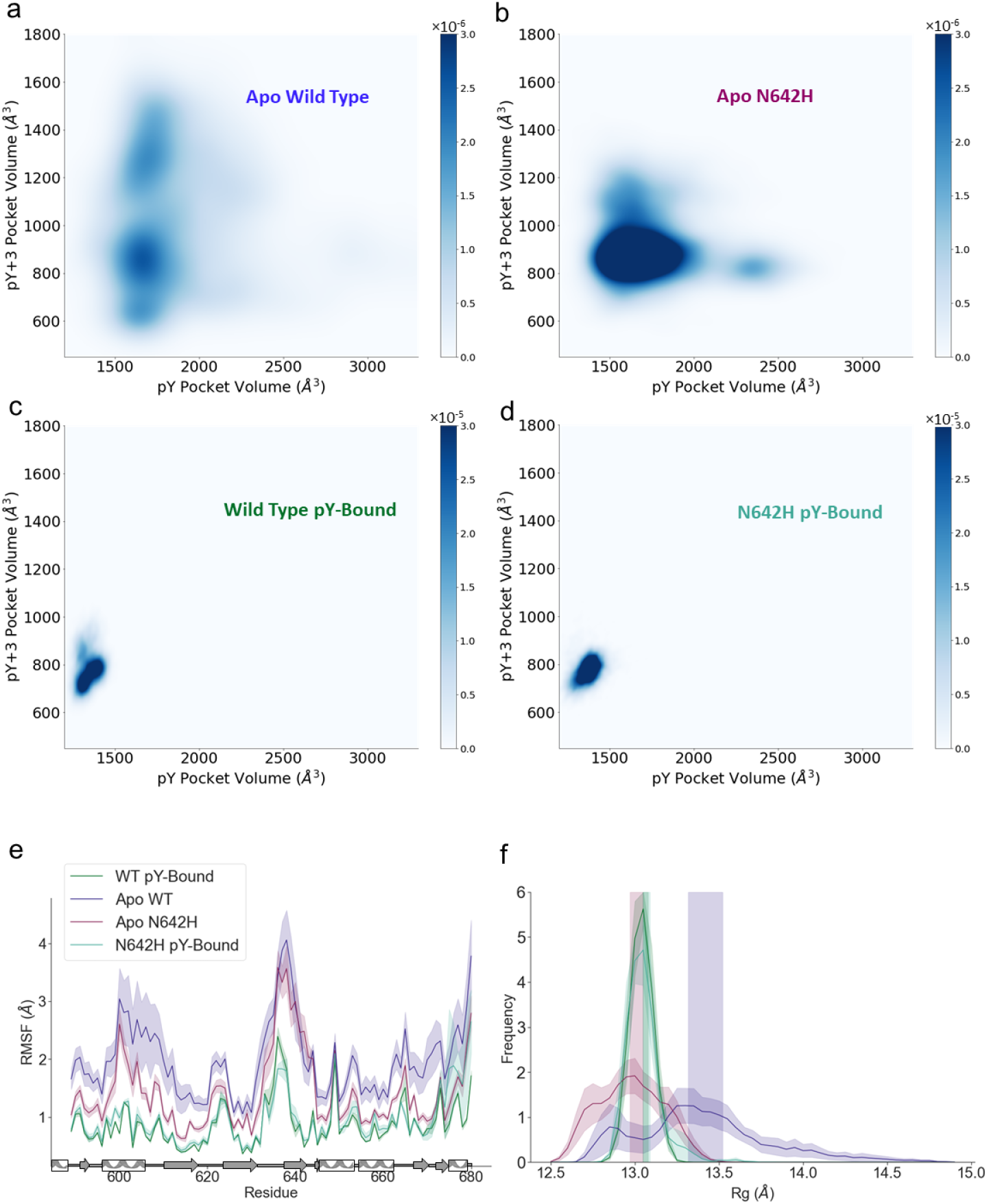
Differences in the structural flexibility of the SH2 domain due to the mutation and parallel dimerization. (a,b,c,d) Two-dimensional distribution of pY pocket volume and pY+3 pocket volume for apo STAT5B (a), apo STAT5B^*N*642*H*^ (b), STAT5B parallel dimer (c), and STAT5B^*N*642*H*^ parallel dimer (d). The colour of each pixel indicates the frequency of sampling each region. Darker colours indicate more frequently sampled regions. Different colour scales are used in panels (a,b) and (c,d). (e,f) Root-mean-square fluctuation (RMSF) profiles (e) and Radius of gyration (*R_g_*) distributions (f) for STAT5B parallel dimer (dark green), STAT5B^*N*642*H*^ parallel dimer (teal), apo STAT5B (blue), and apo STAT5B^*N*642*H*^ (plum) ensembles. The vertical shaded areas indicate the average value of *_Rg_*± the standard error of the mean. In (e) and (f), shaded areas indicate standard error of the mean obtained by treating each SH2 domain in the simulations as an independent measurement.

The rigidifying effect of the N642H mutation occurs not only in the pY+3 pocket, but also globally. The root-mean-square fluctuation (RMSF) profile (Figure 3e) shows a decreased flexibility through-out the SH2 domain with the mutation. The most flexible region in the SH2 domain includes the CD loop and the *β*D-strand (residues 630-645). While this region contains the mutation site, it shows no difference in flexibility between apo STAT5B and apo STAB5B^*N*642*H*^. The reduced flexibility of STAT5B^*N*642*H*^ compared to wild-type STAT5B (Figure 3e) is also seen in the crystallographic B-factors (Figure S12) reported for the structures of the STAT5B and STAB5B^*N*642*H*^ antiparallel dimers ^9^. Compared to either of the apo systems, the parallel dimer has lower RMSF (Figure 3e), indicating even greater rigidity. This is further indicated by stabilization of all *α*-helices and the *β*-sheet in the parallel dimer (Figure S8b,c). In general, SH2 domains are known to be conformationally flexible ^36;61^. The increase in rigidity in the parallel dimer that we observe is consistent with experimental studies that reported increased order of the pY+3 pocket ^55^ and global stabilization of the SH2 domain ^37^ upon ligand binding.

Rigidification of the SH2 domain due to the N642H mutation can be seen in other structural properties, including R*_g_* (Figure 3f), which is more narrowly distributed for STAT5B^*N*642*H*^ compared to STAT5B. It is also seen in the 2D contact difference map of apo STAT5B^*N*642*H*^ and apo STAT5B (Figure S8d), which shows that more intramolecular contacts are formed on average in STAT5B^*N*642*H*^. To further understand the differences in the conformational spaces explored by the simulations, we performed PCA on the simulation trajectories (Figure S8e). Based on the projection of the trajectories in this conformational space, we observe that the apo wild-type simulations exhibit a significantly more diverse conformational ensemble, consistent with greater flexibility. In fact, the results of PCA demonstrate that both pY binding and the N642H mutation have a rigidifying influence on the SH2 domain globally.

Taken together, these results indicate that the N642H mutation and pY binding have a similar influence on the conformational dynamics of the SH2 domain. Both apo STAT5B^*N*642*H*^ and the STAT5B parallel dimer are more rigid than apo STAT5B, as shown by several metrics. These differences are visible in the structural ensembles of these systems (Figure S13a-f). The reduction in the flexibility of the SH2 domain caused by the N642H mutation could contribute to the increased parallel dimerization affinity of the protein by offsetting the entropic cost of pY binding, which, as demonstrated above, also rigidifies the protein.

### The N642H mutation stabilizes a parallel-dimer-like conformation

In the crystal structure of the STAT5B^*N*642*H*^ antiparallel dimer, the SH2 domain of each protomer is in a different conformational state ^9^. In one of these states, which we call the A-state, the *β*D-strand is dissociated from the rest of the *β*-sheet. In the other state, which we call the B-state, the 3-stranded *β*-sheet is well-formed. Our simulations, which use this crystal structure as a starting point, show that these two states persist in simulation, which is visible in the structural ensembles taken from the simulations (Figure S13g,h). Furthermore, a comparison between the parallel dimer and the STAT5B^*N*642*H*^ antiparallel dimer ensembles reveals that the structure of the B-state is highly similar to the structure of the SH2 domain of the parallel dimers (Figure S13c,f,h).

To assess the similarity between the B-state and the structure of the SH2 domain observed in the parallel dimer simulations, we carried out a principal component analysis on the STAT5B^*N*642*H*^ antiparallel dimer and parallel dimer systems. This analysis (Figure 4d-g) shows that the SH2 domain of the STAT5B^*N*642*H*^ antiparallel dimer samples multiple states, which can be categorized into two macrostates that are similar to the A-state and B-state in the crystal structure of the STAT5B^*N*642*H*^ antiparallel dimer ^9^. In this projection of the conformational space, the B-state is a conformational state that is intermediate between the A-state and the parallel dimer conformational state. We do not observe transitions between these states in any of the STAT5B^*N*642*H*^ trajectories. Analysis of the secondary structure of these two states (Figure 4b) demonstrates that the A-state lacks a *β*-strand in the region spanning residues 633 to 645. In contrast, this region has a significantly higher propensity to form a *β*-sheet in the B-state (corresponding to the *β*C residues 633-635 and *β*D residues 638-645, Figure 4b), which provides rigidity to the SH2 domain (Figure 4c). The pattern of secondary structure in the STAT5B^*N*642*H*^ B-state ensemble resembles that of the parallel dimer ensembles. Additionally, the mutation site, residue 642, forms similar interactions with the *α*A-helix, *β*B-strand, and *β*C-strand (Figure 4a) in the B-state and pY-bound ensembles. Figure S13c,f,h shows the two distinct states sampled by the STAT5B^*N*642*H*^ antiparallel dimer and the structural similarity between the B-state and pY-bound ensembles. Taken together, these results indicate that the N642H mutation in the SH2 domain stabilizes an active-like, intermediate conformational state, which could contribute to the observed hyperactivation of STAT5B^*N*642*H*^.

**Figure 4:**
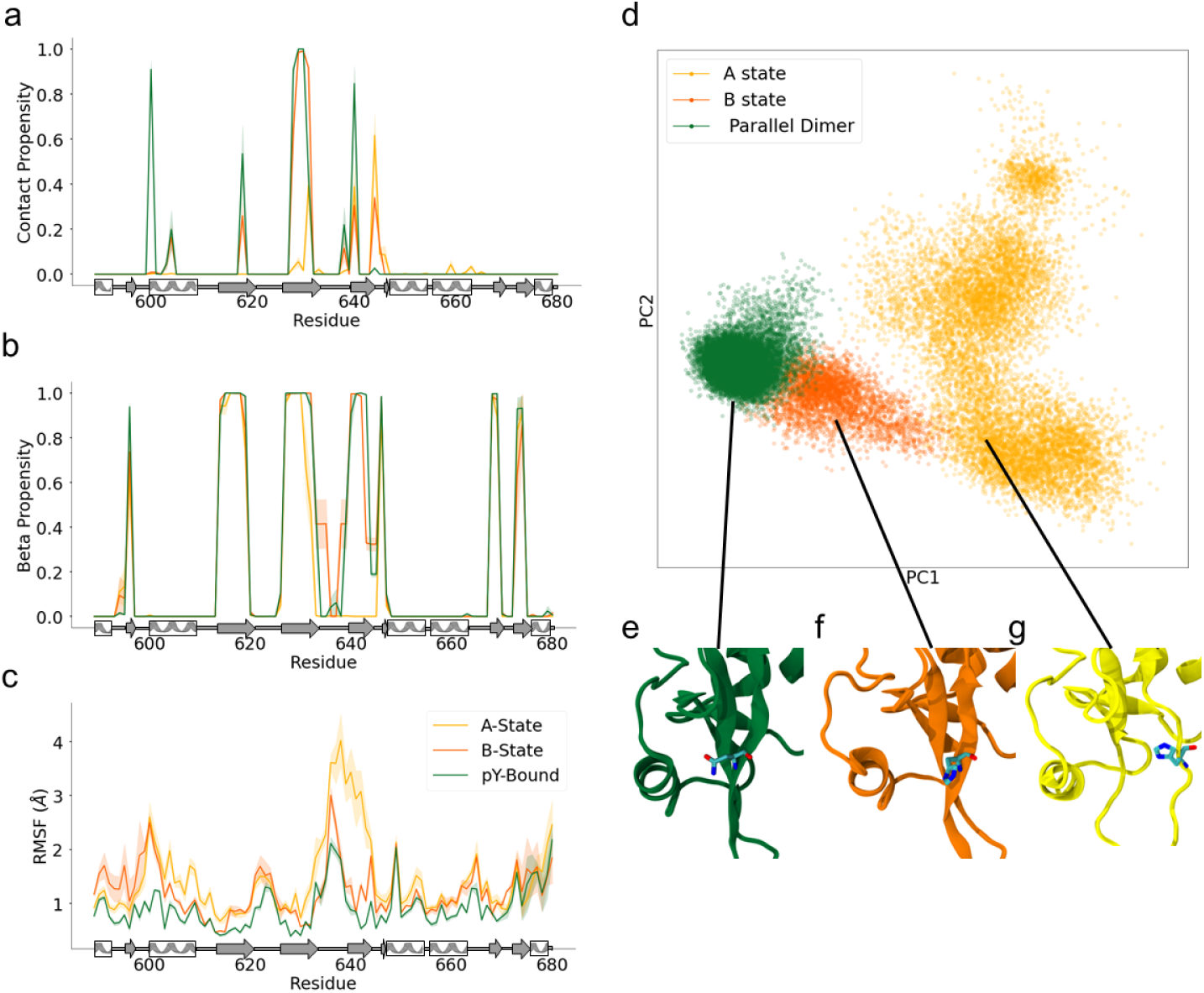
Comparison of N642H mutant antiparallel dimer A-state and B-state trajectories with pY-bound parallel dimer ensemble. (a) Average number of contacts between the mutation site residue (642) and other SH2 domain residues. (b) *β* propensity of each residue in the SH2 domain. (c) Root mean squared fluctuation of each residue in the SH2 domain. The secondary structure is shown below the x-axis. In (a-c), shaded regions indicate standard error of the mean obtained by treating each SH2 domain in the simulations as an independent measurement. (d) Principal component analysis of all N642H mutant antiparallel dimer and both WT and N64H mutant pY-bound parallel dimer trajectories. The B-state ensemble partly overlaps with the parallel dimer ensemble, which is disjoint with the A-state ensemble. (e-g) Close-up view of representative structures from the parallel dimer (e), B-state (f), and A-state (g) ensembles. The mutation site residue (642) is shown in licorice representation. In all figure panels, results for the A-state, B-state and parallel dimer are indicated in yellow, orange and green, respectively. Trajectories are separated into A-state and B-state trajectories based on the occurrence of any *β* structure in the *β*D-strand.

A projection of all STAT5B simulation trajectories onto the STAT SH2 conformational landscape we constructed from crystal structures illustrates the effects of the N642H mutation on the structural ensembles (Figure 5a-f). In this space, the parallel dimer simulations occupy compact, overlapping regions, indicative of their compact, rigid, and broadly similar structure. The apo STAT5B systems widely explore the breadth of this conformational space, indicating a greater degree of conformational heterogeneity compared to the apo STAT5B^*N*642*H*^ simulations, which explore a much more restricted area. The heterogeneous ensemble of apo STAT5B includes some conformations with the *β*-strands that characterize the B-state of STAT5B^*N*642*H*^ (Figure S8c). The A-state and B-state of the STAT5B^*N*642*H*^ antiparallel dimer occupy distinct regions of the space. The B-state overlaps with the same region of the space as the parallel dimer trajectories, indicating their conformational similarity.

**Figure 5:**
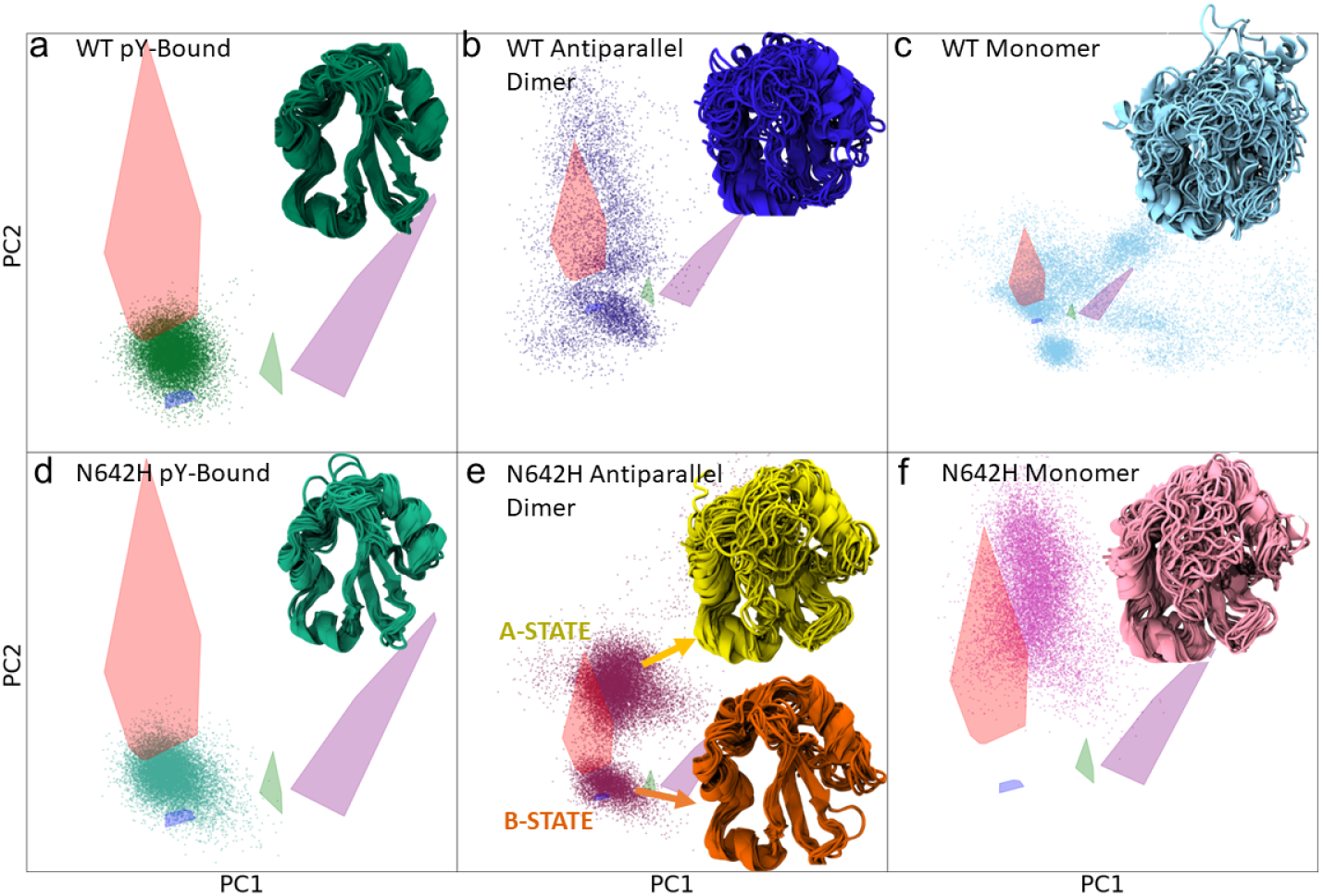
The STAT SH2 Conformational Landscape. (a-f) Every structure from all simulation trajectories was projected onto the conformational space of STAT SH2 domains (the same principal components as in Figure 2a). Each point represents one structure. Structural ensembles consisting of 24 aligned structures selected at random from simulation trajectories are also shown for each system: STAT5B parallel dimer (a), antiparallel dimer (b), and monomer (c); STAT5B^*N*642*H*^ parallel dimer (d), antiparallel dimer (e), and monomer (f).

In this space, the apo STAT5B ensemble includes conformations found in the same regions as the STAT5B^*N*642*H*^A-state and B-state, suggesting that apo STAT5B may access these states as well. To better understand the conformational ensemble of apo STAT5B in terms of the A-state and B-state, we projected the apo STAT5B trajectories onto the PCA space from Figure 4d, which was obtained using only the parallel dimer and STAT5B^*N*642*H*^ antiparallel dimer trajectories (Figure S14). This projection shows that the STAT5B monomer and antiparallel dimer ensembles both overlap with the A-state and B-state regions of this space. However, the A-state and B-state are not as clearly delineated in the apo STAT5B ensemble. These results suggest that conformations similar to the A-state and B-state exist within the heterogeneous ensembles of the STAT5B monomer and antiparallel dimer.

To see if other STAT proteins exhibit conformational heterogeneity in their SH2 domains, we carried out additional simulations of the SH2 domains of STAT1 and STAT5A. We then projected the resulting structural ensembles onto the STAT SH2 conformational landscape (Figure S15a, Figure S16a). Both of these systems display a state with a dissociated *β*D-stand and a state with a well-formed *β*-sheet. These states arise spontaneously, although the crystal structures from which the simulations were initiated (1YVL ^62^ and 1Y1U ^63^) both have a well-formed *β*-sheet. While we do not observe transitions between the A-state and B-state in the STAT5B^*N*642*H*^ simulations, we do observe transitions between the two states in the simulations of both STAT1 and STAT5a (Figure S15b, Figure S16b). These observations of the *β*D-strand sampling two conformational states suggest this may be a common feature among STAT SH2 domains. The clear delineation of these states in the STAT5B^*N*642*H*^ ensemble, but not the STAT5B ensemble, suggests that the N642H mutation stabilizes these distinct conformational states of the SH2 domain.

### Hydrogen deuterium exchange experiments show similar impact of the N642H mutation and pY binding

To validate the results of our simulations showing an effect of the N642H mutation on the structure and dynamics, we carried out millisecond time-resolved hydrogen deuterium exchange mass spectrometry (TRESI-HDX) ^64;65^. This approach uses the rate of exchange of amide protons from the peptide backbone with deuterium from solvent to measure the relative stability of the hydrogen bonding network that maintains secondary and tertiary structures in proteins. In general, dynamic regions of proteins exhibit faster deuterium uptake compared to structured regions of proteins. When conducted on the millisecond timescale, these measurements are sensitive even to subtle shifts in conformational dynamics resulting from mutation and/or ligand binding.

First, we compare the difference in deuterium uptake between the phosphopeptide bound and unbound STAT5B (Figure 6a,b). The HDX results show decreased deuterium uptake rates in the SH2 domain upon phosphopeptide binding, suggesting a stabilization of secondary and tertiary structures in the pY-bound state. Next, we determine the difference in deuterium uptake between apo STAT5B and STAT5B^*N*642*H*^ (Figure 6c,d). The results of this HDX analysis show decreased deuterium uptake in STAT5B^*N*642*H*^ compared to wild type. This decreased deuterium uptake indicates that the N642H mutation results in strong stabilization of secondary and tertiary structures in the immediate vicinity. Interestingly, the effect of the mutation is highly confined, with no evidence of significant changes to dynamics ortho-or allosteric to the mutation site. Together, these data are in support of our computational predictions, indicating that both phosphopeptide binding and N642H mutation have a similar stabilizing influence on the SH2 domain.

**Figure 6:**
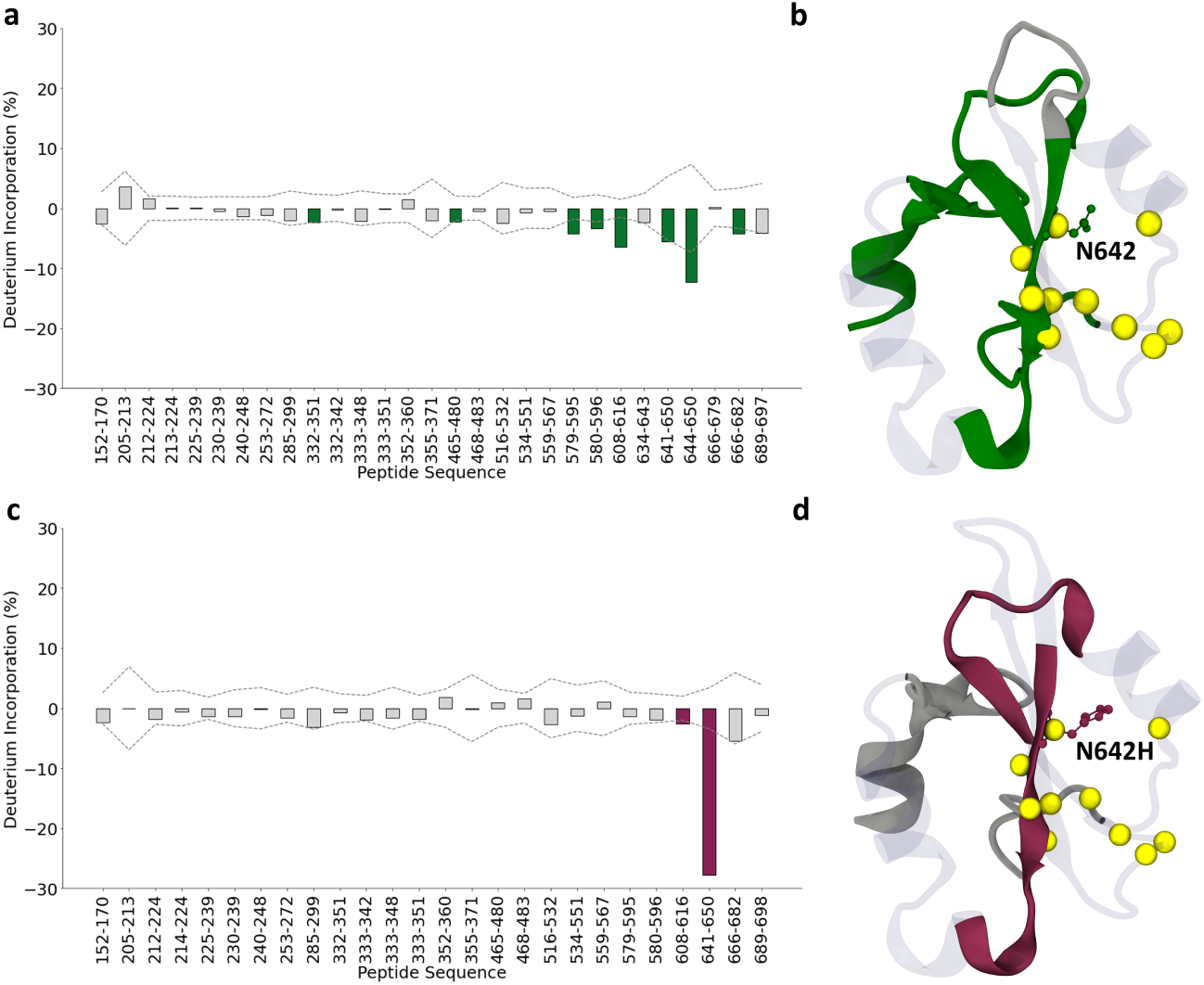
HDX experiments show a difference in deuterium uptake upon pY binding and with the N642H mutation. (a) The difference in deuterium uptake between apo STAT5B and phosphopeptide-bound STAT5B. Green bars indicate peptides having significantly greater deuterium uptake in the apo case. Uncertainty is indicated by dashed lines in panels (a) and (c). (b) A structure of the SH2 domain taken from the first frame of the wild-type parallel dimer simulation. Residues belonging to peptide sequences that show greater deuterium incorporation in the apo state relative to pY bound are shown in green. In both panels (b) and (d), residues belonging to peptide sequences that show no difference in deuterium incorporation are shown in grey, regions for which there is no deuterium incorporation data are shown as transparent, C*α* atoms of pocket residues are shown as yellow spheres, and the N642 residue is shown in a ball and stick representation. (c) The difference in deuterium uptake between apo STAT5B and apo STAT5B^*N*642*H*^. Plum bars indicate peptides having significantly greater deuterium uptake in wild type. (d) A structure of the SH2 domain taken from the first frame of the N642H parallel dimer simulation. Peptide sequences that show decreased deuterium incorporation in the N642H mutant are shown in plum.

## Discussion

This work provides a detailed first description of the parallel dimer structure of the STAT5B SH2 domain and how it is influenced by the N642H mutation. Our simulations reveal multiple N642H-induced changes that could contribute to the hyperactivity and drug resistance of STAT5B^*N*642*H*^. We find a hydrogen bonding path initiating directly from N642H that leads to strengthened hydrogen bonds between the phosphotyrosine and R618, the most highly conserved residue across all SH2 domains. We also find that the pY residue has increased interactions with the N642H residue itself as well as other residues near the mutation site, which further strengthen the binding interaction. Furthermore, we find that the mutation rigidifies the structure of the apo-SH2 system, thereby reducing the conformational entropy. At the same time, we observe that the pY-bound systems are extremely rigid, indicating an entropy decrease upon formation of the parallel dimer. Thus, the entropic cost required for binding is smaller for the mutant, making parallel dimerization more favourable. We have not identified any simple mechanism for this effect of the N642H mutation; it likely arises from the change in probability of multiple interactions that together stabilize the central *β*-sheet of the SH2 domain. Finally, we show that the STAT5B^*N*642*H*^ accesses a conformational state that closely resembles the phosphorylated parallel dimer conformation, even in the unphosphorylated state. Stabilization of a parallel-dimer-like conformation even in the absence of phosphopeptide could suggest increased stability of the conformation in the presence of phosphopeptide, which may lead to prolonged parallel dimerization. Together, the combination of these disparate N642H-induced structural and dynamic changes plausibly leads to the increased parallel dimer affinity associated with the hyperactivity and oncogenicity of STAT5B^*N*642*H*^. While it is possible that all of these effects contribute to the oncogenicity of STAT5B^*N*642*H*^, we consider the enhanced pY binding observed in the parallel dimer to be the most likely contributor given the wealth of evidence that the pY-R618 interaction is the most critical aspect of SH2 binding.

Our homology model of the STAT5B parallel dimer also allows us to speculate on the structural cause of multiple leukemia-associated mutations in the SH2 domain: T628S, N642H, Y665F, P702A, I704L, and Q706L ^23;66^ (Figure 7). Y665 is a constituent of the hydrophobic pY+3 pocket and directly interacts with the pY+3 residue (Figure S9). The mutation Y665F may increase the hydrophobicity of the pY+3 pocket, leading to enhanced binding and prolonged parallel dimerization. Similarly, Q706 is frequently in contact with the pY+3 residue in the STAT5B parallel dimer simulations. The Q706L mutation could enhance parallel dimerization through hydrophobic interactions between pY+3 and Q706L. Furthermore, Q706 participates in the inter-protomer *β*-sheet and is located side-by-side with a hydrophobic residue on the conjugate protomer (Figure 7b). The increased hydrophobicity of Q706L could, therefore, stabilize the *β*-sheet via enhanced hydrophobic interactions. T628 is a primary constituent of the pY binding pocket, and the T628S mutation is likely to influence dimerization affinity via changes in interactions with the pY residue. Furthermore, T628 is an important hydrogen bonding partner of R618, and the T628S mutation could also influence parallel dimerization affinity via repositioning of R618. Threonine is present at this position in all human STAT proteins ^10^ and all mammalian STAT proteins for which a crystal structure is available (Figure S17). Residue P702 (pY+3) is crucial for determining specificity and is an important constituent of the parallel dimerization interface. The P702A mutation could influence parallel dimer affinity by the alanine packing more efficiently into the pY+3 pocket compared to proline. We also note that an alanine residue is present in this position in the sequence of STAT6 ^10^. Based on the crystal structure of the parallel dimer of STAT6 ^10^, that alanine residue is part of a *β*-strand and fits well into the pY+3 pocket. Changing alanine to proline would likely destabilize this *β*-strand, thereby destabilizing the parallel dimer interface. Similarly, the I704L mutation could influence parallel dimerization by enhancing interactions in the inter-protomer *β*-sheet. STATs 1 and 2 have a leucine at this position ^10^.

**Figure 7:**
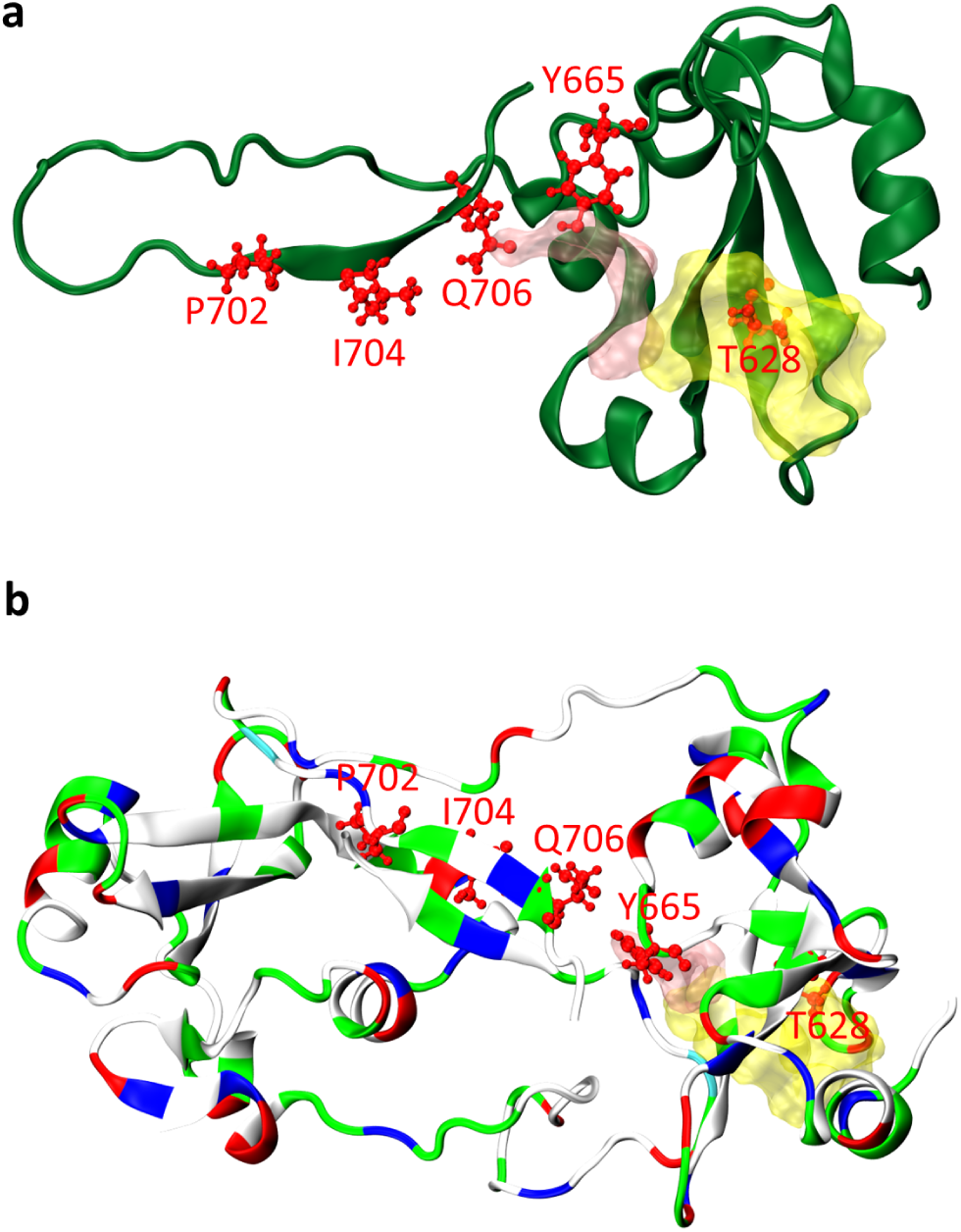
Common cancer mutations in the STAT5B SH2 domain. (a)The STAT5B SH2 domain and is shown in cartoon representation. The pY and pY+3 pockets are shown in surface representation, in yellow and pink, respectively. Sites of mutations that commonly occur in cancer ^66^ are shown in ball and stick representation coloured in red. Residue N642 is also a mutation site, but it is excluded for clarity. Mutations identified at these sites include: T628S, Y665F, P685R, P702A, and Q706L ^66^. (b) The parallel dimer of the STAT5B SH2 domain is shown in cartoon representation, coloured by residue type. Negative residues are shown in red, positive residues are shown in blue, polar residues are shown in green, and hydrophobic residues are shown in white. Sites of mutations that commonly occur in cancer are coloured red and shown in ball-and-stick representation on one of the monomers. The pY and pY+3 are shown in surface representation and coloured in yellow, and pink, respectively.

On the basis of our findings, we speculate that the drug resistance observed in STAT5B^*N*642*H*^ ^28^ may be due to the compounded effect of stronger pY binding and reduced entropic cost allowing parallel dimerization to out-compete binding with small molecule inhibitors that target wild-type STAT5B. Ideally, a novel therapeutic would preferentially bind STAT5B^*N*642*H*^ over the wild type. The structural information presented here could be used to accelerate the rational design of therapeutics targeting STAT5B^*N*642*H*^.

## Methods

### MD simulation methods

For all MD simulations in this study, the CHARMM36m force field ^67^ was used with the CHARMM-modified TIP3P water model ^68^. The N- and C-termini were simulated in the charged state and all titratable residues were simulated in their standard states for pH 7. Periodic boundary conditions were used. Neutralizing Na^+^ and Cl*^−^* ions were added at a concentration of 150 mM. Energy minimization was performed for each system using the steepest descent method. For the initial equilibration simulations, velocities were generated from a Maxwell distribution. The bonds involving hydrogen atoms were constrained using the LINCS algorithm ^69^. The SETTLE algorithm ^70^ was used to constrain the water molecules. Parrinello-Rahman pressure coupling ^71^ and the velocity-rescale thermostat ^72^ were used for all production simulations. Simulations were run at a temperature of 298 K and a pressure of 1 bar. Additional simulation details specific to each system are provided below. Refer to Table 1 for a list of simulation systems and Table S1 for a detailed list of all trajectories analyzed.

### Obtaining a Homology Model for the Parallel Dimer

We constructed a homology model for the STAT5B parallel dimer using the structure of the STAT6 parallel dimer (PDB ID: 4Y5U ^10^) as a template. The resulting homology model, obtained using SWISSMODEL ^73^, had an overall MolProbity ^74^ score of 2.39, with 91.7% of the residues having backbone dihedral angles in the most favoured regions of the Ramachandran plot. Each monomer was modelled separately and the two monomers were then combined to produce the model for the parallel dimer. To avoid steric clashes between the two monomers, it was necessary to perform loop refinement on the region including residues 686-695 using the Modeller ^75^ plugin for UCSF Chimera ^76^. Residue Y699 on both monomers was phosphorylated using the CHARMM-GUI ^77^ PDB reader tool ^78^. After constructing the model of the parallel dimer, we removed all STAT5B domains excluding the SH2 and linker domains from the parallel dimer structure such that each monomer comprised residues 498-709. The number of residues in the linker domain in the homology model is the same as that in the STAT6 template. The SH2 and linker domains of the model and template (STAT6) had a sequence identity of 53.58%, and the SH2 domain specifically had a sequence identity of 52%. See Figure S1b for the sequence of the SH2 domain and phosphotyrosine motif region that was used.

### Simulation Procedure for the Parallel Dimers

After the initial construction of the homology model, the resulting structure was placed in a dodecahedral box with a 1 nm distance to all box edges, adding water molecules and neutralizing Na^+^ and Cl*^−^* ions. Simulations of the parallel dimers were performed using GROMACS version 2019.1 ^79^. Following energy minimization, equilibration simulations were performed with position restraints on all protein heavy atoms, including (1) a 20 ns simulation in the NVT ensemble, (2) a 20 ns simulation in the NPT ensemble using the Berendsen method for pressure coupling ^80^, and (3) a 20 ns in the NPT ensemble with Parrinello-Rahman pressure coupling ^71^. Position restraints were released from residues in the disordered regions (634-638 and 680-695) before being released from the rest of the protein. Position restraints were reduced gradually over the course of four separate 20 ns simulations, for a total of 80 ns, with descending position restraint force constants of 1000 kJ/mol, 100 kJ/mol, 10 kJ/mol, and 1 kJ/mol while keeping force constants of 1000 kJ/mol on all other protein heavy atoms. Following these steps, position restraints were released on all other heavy protein atoms over the course of 80 ns, following the same procedure. Finally, a 20 ns simulation without any position restraints was run before beginning production runs. Production runs comprised 3 independent 2 *µ*s simulations in the NPT ensemble, totalling 12 *µ*s of sampling for a single chain.

The initial structure for the parallel dimer with the N642H mutation was obtained by selecting a random frame from the wild-type parallel dimer production run, extracting the protein coordinates, and introducing the N642H mutation using PyMOL ^81^. The histidine residue at position 642 was simulated in the deprotonated (neutral) state, as PROPKA ^82^ predicted a pK*_a_* value between 2.5-3.25. We also chose to simulate this histidine residue as neutral to be consistent with NMR measurements of two other SH2 domains with a histidine residue in the same position with measured pK*_a_* values below 4.0 ^55;56^. The rest of the simulation procedure is the same as that of the wild-type parallel dimer, but with additional steps to gradually release the position restraints to prevent dimer dissociation due to the perturbation of the interaction between residue 642 and the phosphotyrosine residue. Position restraints on all heavy atoms were gradually released over 9 separate 20 ns simulations, totalling 180 ns, with descending position restraint force constants of 1000 kJ/mol, 500 kJ/mol, 250 kJ/mol, and 100 kJ/mol, 50 kJ/mol, 25 kJ/mol, 10 kJ/mol, 5 kJ/mol, and 1 kJ/mol followed by 20 ns of unrestrained simulation before production simulations were started.

For all parallel dimer simulations, the bonds involving hydrogen atoms were constrained using the LINCS algorithm ^69^, and water molecules were constrained using the SETTLE algorithm ^70^. A timestep of 2 fs was used. A cut-off of 1 nm was used to calculate the short range electrostatic and van der Waals interactions. The non-bonded pair list was updated every 100 fs to include atoms less than 1.1 nm apart. Long range electrostatic interactions were computed using particle mesh Ewald (PME) summation ^83^ with a grid spacing of 0.16 nm and fourth order cubic interpolation. All other simulation details are as described above in *MD simulation methods*.

### Simulation Procedure for the Monomers and Antiparallel Dimers

The simulations of the monomer and antiparallel dimer forms of STAT5B and STAT5B^*N*642*H*^ follow the same procedure as described in de Araujo *et al.* ^9^. The simulations of the monomers were extended to 2 *µ*s, and three additional independent trajectories were added for each system for a total of 12 *µ*s of sampling time per system. Since the simulations of the STAT5B antiparallel dimer from de Araujo *et al.* were terminated due to dimer dissociation, they were not extended. Simulations of the STAT5B^*N*642*H*^ antiparallel dimer were extended to 2 *µ*s. Three additional simulations of both the STAT5B and STAT5B^*N*642*H*^ antiparallel dimer were performed. These additional simulations of STAT5B antiparallel dimers dissociated shortly after 1 *µ*s and the trajectories were truncated at this point (specifically, at 1.144 *µ*s). In total, we analyzed 1.144 *µ*s *×* 2 chains *×* 3 trajectories = 6.86 *µ*s of single-chain samplng of the STAT5B antiparallel dimer. The three additional simulations of STAT5B^*N*642*H*^ antiparallel dimers also showed dissociation. One dissociated at the beginning of the simulation and was therefore not included in the analysis. The other two dissociated at 1.228 *µ*s and 1.218 *µ*s, respectively. In total, we analyzed 16.9 *µ*s of single-chain sampling (2 *µ*s *×* 2 chains *×* 3 trajectories + 1.228 *µ*s *×* 2 chains *×* 1 trajectory + 1.218 *µ*s *×* 2 chains *×* 1 trajectory = 16.9 *µ*s) for this system.

Only a small number of contacts are present between the protomers in both the STAT5B and STAT5B^*N*642*H*^ antiparallel dimer crystal structures, which may explain why they dissociate rapidly in our simulations ^22^. Furthermore, the N-terminal domain, which is critical for the formation of unphosphorylated STAT5B dimers in cells ^18^, is missing in the crystal structures and in our simulations. This likely also contributes to the observed instability of the antiparallel dimer structure. In addition, for unphosphorylated STAT5A and the unphosphorylated core fragment of STAT3, the monomer is known to be the dominant state in solution ^13;17^. Similarly, STAT5B may prefer the monomeric state, which could also explain the fast dissociation observed in the simulations of the antiparallel dimers.

The simulations of STAT1 and STAT5A were initiated from crystal structures (STAT1 PDB: 1YVL ^62^ chain A, STAT5A PDB: 1Y1U ^63^ chain A). Only the linker and SH2 domains were simulated (STAT1: L488-S682, STAT5A: H471-A690). The missing atoms in the crystal structure of STAT1 were added using Swiss-PDB Viewer ^84^ and the missing loops were introduced using the loop model module of Modeller ^75^. The resulting loop models were refined using the refine fast option of loop model. Six different structures for the loop regions were obtained as a result of loop modeling and refinement, which were used as initial structures in six independent simulations for each dimer. Six independent simulations were performed for each protein. These simulations followed the same procedure as described in de Araujo *et al.* ^9^.

### Defining the Binding Pockets

Residues comprising the pY and pY+3 binding pockets in the SH2 domain were determined based on the contacts formed by these residues (pY699 and P702) in the simulations of the wild-type and N642H parallel dimers. Both chains in all parallel dimer simulations (a total of 24 *µ*s of sampling) were analyzed to compute the inter-chain contacts formed with these residues. A contact between two residues is considered to be present if any pair of atoms is within a distance cut-off of 3.5 Å. Only the contacts occurring in at least a tenth of the simulation frames in both the N642H and wild-type parallel dimer simulations were included in the definitions of the pY and pY+3 pockets. Using this approach to define the binding pockets, the residues comprising the pY pocket were found to be consistent between the N642H and wild-type simulations. The pY binding pocket comprises residues K600, R618, S620, D621, S622, T628, N642/H642, L643, M644, and P645 (Figure S5). The same methodology applied to the pY+3 binding pocket results in significant variation between the wild-type and N642H systems due to differences in the persistence of contacts formed between the pY+3 residue and residues 706-709 (Figure S9b,c). Residues 706-709 are not present in the monomer and antiparallel dimer systems, and were therefore excluded from the pY+3 pocket definition for all analysis. As a result, the pY+3 pocket comprises residues L643, M644, F646, L663, and Y665. We note that residues 706-709 were not included in the constructs used to obtain the crystal structures ^9^ used as the initial structures of the antiparallel dimer simulations, so we did not include these residues in those simulations.

### Constructing the STAT SH2 Domain Conformational Landscape

We performed two-dimensional principal component analysis (PCA) on the set of STAT SH2 domain crystal structures that were deposited in the Protein Data Bank (PDB) as of August 2024. The PDB accession codes and the chains used for this analysis can be found in Table S2. Similar use of PCA has been reported previously to study the conformational landscapes of proteins ^85–87^.

A feature vector for PCA was chosen using a common set of residues between all structures. A structure-based sequence alignment of the entire set of structures was performed using the MUS-TANG software package ^88^. Only 55 residues aligned between all the structures, due to insertions or deletions between them (Figure S17). We computed the pairwise distance between the C*α* atoms of all 55 aligned residues for each structure. These distances were flattened into a 1D array (length = ½ × 55 × 54 = 1485) representing the conformation of each structure. These feature vectors were used for the PCA. We used the scikit-learn ^89^ toolkit to preprocess the data for PCA, by normalizing and centering each feature. PCA to two dimensions was carried out using the PCA module in scikit-learn ^89^. The first two principal components (PCs) capture 35% and 10% of the variance in the data, respectively.

We plotted the STAT SH2 domain structures in the resulting two-dimensional conformational space. In this two-dimensional conformational space, the structures tend to cluster with other structures of the same STAT protein such that the space can be partitioned into distinct regions containing each STAT protein: STAT1, STAT2, STAT3, STAT5, and STAT6 (Figure S3). The simulation trajectories were then mapped onto this two-dimensional PCA space by constructing a feature vector of pairwise distances for the 55 corresponding residues and applying the pre-computed normalization, centering, and PCA transformation to each trajectory.

The principal axes can be physically interpreted by examining which features have the largest coefficients in the linear expansion of each principal axis. The top 5 contributing residue pairs to the vertical axis suggest that the vertical axis is largely governed by the proximity of *β*D-strand to *β*B-strand and the N-terminus. Overall, we see large contribution to this principal axis from distances between residues (640,645) -(596,620) (Figure S4). This indicates that SH2 domains that are far from each other along the vertical axis have a large difference in the positioning of the *β*D-strand. The horizontal principal axis corresponds largely to the distance between the *α*A-helix and the *α*B’-helix (Figure S4). Similarly, the horizontal axis’ largest contributions suggest that it is largely governed by the distance between the *α*A-helix (and its adjacent loops) and *α*B’-helix (and its N-terminally adjacent loop).

### Analysis of Simulation Trajectories

The simulation trajectories were visualized using VMD ^90^. Residue numbering corresponds to the UNIPROT residue numbers for human STAT5B (UNIPROT ID: P51692). The first 250 ns of each production run was treated as an equilibration period and omitted from analysis. Each protein chain of each independent trajectory is treated as an independent measurement and uncertainty is reported as the standard error of the mean of these measurements. Secondary structure was analyzed with the DSSP ^91^ function in the MDTraj ^92^ module for Python 3. Helix/beta propensity is defined as the portion of frames that a given residue has a helical/beta conformation. Solvent accessible surface area (SASA) was computed using the Shrake-Rupley algorithm ^93^ as implemented in MDTRAJ ^92^ with a probe radius of 1.4 Å and 960 sphere points. All atomic positions, inter-atomic distances, radius of gyration (R*_g_*), root mean square deviation (RMSD), and root mean square fluctuation (RMSF) were computed using the MDAnalysis ^94^ module for Python 3. Two residues were considered to be in contact based on a 3.5 Å cutoff between any pair of atoms. The pocket volume was defined as the convex hull volume of all atoms in all pocket residues. The convex hull was determined with the Open3D ^95^ module for Python 3 after extracting the atomic coordinates using MDAnalysis ^94^. Hydrogen bonds were computed using MDAnalysis with a 3.5 Å distance cutoff between the donor and acceptor atoms and a donor-hydrogen-acceptor angle cutoff of 150 degrees. PCA was performed on normalized and centered C*α* pairwise distances using the scikit-learn ^89^ standard preprocessing library before dimensionality reduction using the sci-kit learn ^89^ PCA module. *π* -*π* stacking interactions were analyzed using the ring centroid distance and the ring stacking angle. The centroid distance between rings was computed as the distance between the mean position of all heavy atoms in each ring. The stacking angle was computed as the smallest angle between the normal vector to each ring. The normal vector to each ring was computed by the following process. For each ring, the displacement vector between every pair of heavy atoms was computed and normalized to unity. Since all of these vectors should lie in the plane of the ring, the cross product of any of them should produce a normal vector to the plane. To reduce errors due to minute fluctuations of the ring planarity, the cross product was taken between every pair of vectors in the ring and the x-positive normal vector was chosen arbitrarily. The orientation vector of each ring was defined as the average of all normal vectors and was normalized to unity. The stacking angle was computed as the arccosine of the dot product of the orientation vector for each ring.

### Experimental Methods

The expression and purification of STAT5B and STAT5B^*N*642*H*^ followed the procedure described in de Araujo et al. ^9^. STAT5B (136-703) and STAT5B^*N*642*H*^ (136-703) samples were prepared for time-resolved electrospray ionization mass spectrometry (TRESI-MS) by buffer exchange into 100 mM ammonium acetate (pH 7.0) using Thermo Scientific Zeba Spin Desalting Columns (7K MWCO). 10 *µ*M of the phosphorylated peptide (Ac-QD*pY* LVLDKWL), dissolved in 100% DMSO, was incubated with 5 *µ*M of STAT5B and STAT5B^*N*642*H*^ for 1 hour on ice. TRESI-HDX was performed on Waters Synapt G2-S using microfluidic set up for HDX experiment as previously described ^96;97^. PerkinElmer Series 200 pumps and autosamplers were used to inject protein and deuterium at 5 *µ*l/min over three reaction time points (0.37s, 0.678s, and 15s). Deuterated protein was first quenched with 3% formic acid and 40 mM ammonium acetate (0°C, 40*µ*l/min), then passed through an online digest column made of pepsin linked agarose beads in a peek column with an ID of 0.040” and a 2*µ*m pore size upstream of the source ^97^. Peptic peptides were identified using MS/MS and their deuterium uptake levels were calculated using Mass Spec Studio ^98^. The differences in deuterium uptake between apo and peptide-bound STAT5B and STAT5B^*N*642*H*^ were calculated, and the summed deuterium uptakes for all time points were reported. Reported values are an average of three technical replicates, and the uncertainty is reported as the standard deviation.

## Author Contributions

**Liam Haas-Neill**: -Conceptualization, Formal Analysis, Investigation, Methodology, Software, Visualization, Writing -Original Draft Preparation; **Deniz Meneksedag-Erol**: Conceptualization, Formal Analysis, Investigation, Methodology, Writing -Original Draft Preparation; **Ayesha Chaudhry**: Conceptualization, Formal Analysis, Investigation, Methodology, Writing -Original Draft Preparation; **Masha Novoselova**: Formal Analysis, Investigation; **Qirat Ashraf** : Investigation, Resources; **Elvin de Araujo**: Investigation, Resources; **Derek Wilson**: Conceptualization, Funding Acquisition, Project Administration, Supervision, Writing -Review & Editing; **Sarah Rauscher**: Conceptualization, Formal Analysis, Funding Acquisition, Project Administration, Supervision, Writing -Review & Editing.

## Acknowledgments

This research was supported by a Connaught New Researcher Award to S.R. and a Natural Science and Engineering Research Council of Canada (NSERC) Discovery Grant (RGPIN-201806408). This research was enabled in part by support provided by Calcul Quebec (www.calculquebec.ca) and the Digital Research Alliance of Canada (alliance.can.ca). This research was also supported by an NSERC Discovery Grant and Collaborative Research and Development Grant (RGPIN-480432 and CRDPJ-507056) to D. J. W.

## Data Availability

MD simulation data (initial coordinates and simulation trajectories) are provided for all systems as a Zenodo repository accessible here: https://doi.org/10.5281/zenodo.13942609. Data underlying figures are also available in this Zenodo repository.

## Notes

https://doi.org/10.5281/zenodo.13942609

### Competing Interest Statement

The authors have declared no competing interest.

### Summary of Updates

We added an additional supporting figure and made minor revisions to improve clarity.

## References

1. Li YJ, Zhang C, Martincuks A, Herrmann A, Yu H (2023) STAT proteins in cancer: orchestration of metabolism. Nature Reviews Cancer 23:115–134.

2. Furqan M, Akinleye A, Mukhi N, Mittal V, Chen Y, Liu D (2013) STAT inhibitors for cancer therapy. Journal of Hematology & Oncology 6:90.

3. Hu X, li J, Fu M, Zhao X, Wang W (2021) The JAK/STAT signaling pathway: from bench to clinic. Signal Transduction and Targeted Therapy 6:402.

4. Xue C, Yao Q, Gu X, Shi Q, Yuan X, Chu Q, Bao Z, Lu J, Li L (2023) Evolving cognition of the JAK-STAT signaling pathway: autoimmune disorders and cancer. Signal Transduction and Targeted Therapy 8:204.

5. Philips RL, Wang Y, Cheon H, Kanno Y, Gadina M, Sartorelli V, Horvath CM, Darnell JE, Stark GR, O’Shea JJ (2022) The JAK-STAT pathway at 30: Much learned, much more to do. Cell 185(21):3857–3876.

6. Liu X, Robinson GW, Gouilleux F, Groner B, Hennighausen L (1995) Cloning and expression of STAT5 and an additional homologue (STAT5b) involved in prolactin signal transduction in mouse mammary tissue. Proceedings of the National Academy of Sciences 92(19):8831–8835.

7. Buitenhuis M, Coffer PJ, Koenderman L (2004) Signal transducer and activator of transcription 5 (STAT5). The International Journal of Biochemistry & Cell Biology 36(11):2120–2124.

8. Verhoeven Y, Tilborghs S, Jacobs J, De Waele J, Quatannens D, Deben C, Prenen H, Pauwels P, Trinh XB, Wouters A, Smits EL, Lardon F, van Dam PA (2020) The potential and controversy of targeting STAT family members in cancer. Seminars in Cancer Biology 60:41–56.

9. de Araujo ED, Erdogan F, Neubauer HA, Meneksedag-Erol D, Manaswiyoungkul P, Eram MS, Seo HS, Qadree AK, Israelian J, Orlova A, Suske T, Pham HTT, Boersma A, Tangermann S, Kenner L, Rülicke T, Dong A, Ravichandran M, Brown PJ, Audette GF, Rauscher S, Dhe-Paganon S, Moriggl R, Gunning PT (2019) Structural and functional consequences of the STAT5B N642H driver mutation. Nature Communications 10:2517.

10. Li J, Rodriguez JP, Niu F, Pu M, Wang J, Hung LW, Shao Q, Zhu Y, Ding W, Liu Y, Da Y, Yao Z, Yang J, Zhao Y, Wei GH, Cheng G, Liu ZJ, Ouyang S (2016) Structural basis for DNA recognition by STAT6. Proceedings of the National Academy of Sciences 113:13015–13020.

11. Levy DE, Darnell JE (2002) Stats: transcriptional control and biological impact. Nature Reviews Molecular Cell Biology 3(9):651–662.

12. Awasthi N, Liongue C, Ward AC (2021) STAT proteins: a kaleidoscope of canonical and non-canonical functions in immunity and cancer. Journal of Hematology & Oncology 14:198.

13. Bernadó P, Pérez Y, Blobel J, Ferńandez-Recio J, Svergun DI, Pons M (2009) Structural characterization of unphosphorylated STAT5a oligomerization equilibrium in solution by small-angle X-ray scattering. Protein Science 18(4):716–726.

14. Rawlings JS, Rosler KM, Harrison DA (2004) The JAK/STAT signaling pathway. Journal of Cell Science 117:1281–1283.

15. Park HJ, Li J, Hannah R, Biddie S, Leal-Cervantes AI, Kirschner K, Flores Santa Cruz D, Sexl V, Göttgens B, Green AR (2015) Cytokine-induced megakaryocytic differentiation is regulated by genome-wide loss of a uSTAT transcriptional program. The EMBO Journal 35(6):580–594.

16. Decker T (2016) Emancipation from transcriptional latency: unphosphorylated STAT5 as guardian of hematopoietic differentiation. The EMBO Journal 35(6):555–557.

17. Ren Z, Mao X, Mertens C, Krishnaraj R, Qin J, Mandal PK, Romanowski MJ, McMurray JS, Chen X (2008) Crystal structure of unphosphorylated STAT3 core fragment. Biochemical and Biophysical Research Communications 374(1):1–5.

18. Begitt A, Krause S, Cavey JR, Vinkemeier DE, Vinkemeier U (2023) A family-wide assessment of latent STAT transcription factor interactions reveals divergent dimer repertoires. Journal of Biological Chemistry 299(5):104703.

19. Lin JX, Li P, Liu D, Jin HT, He J, Rasheed MAU, Rochman Y, Wang L, Cui K, Liu C, Kelsall BL, Ahmed R, Leonard WJ (2012) Critical role of STAT5 transcription factor tetramerization for cytokine responses and normal immune function. Immunity 36(4):586–599.

20. Emenecker RJ, Griffith D, Holehouse AS (2021) Metapredict: a fast, accurate, and easy-to-use predictor of consensus disorder and structure. Biophysical Journal 120(2):4312–4319.

21. Emenecker RJ, Griffith D, Holehouse AS (2022) Metapredict v2: An update to metapredict, a fast, accurate, and easy-to-use predictor of consensus disorder and structure. bioRxiv p. 10.1101/2022.06.06.494887.

22. de Araujo ED, Orlova A, Neubauer HA, Bajusz D, Seo HS, Dhe-Paganon S, Keserű GM, Moriggl R, Gunning PT (2019) Structural implications of STAT3 and STAT5 SH2 domain mutations. Cancers 11(11):1757.

23. Küçük C, Jiang B, Hu X, Zhang W, Chan JKC, Xiao W, Lack N, Alkan C, Williams JC, Avery KN, Kavak P, Scuto A, Sen E, Gaulard P, Staudt L, Iqbal J, Zhang W, Cornish A, Gong Q, Yang Q, Sun H, d’Amore F, Leppä S, Liu W, Fu K, de Leval L, McKeithan T, Chan WC (2015) Activating mutations of STAT5b and STAT3 in lymphomas derived from *γδ*-t or NK cells. Nature Communications 6:6025.

24. Klein K, Kollmann S, Hiesinger A, List J, Kendler J, Klampfl T, Rhandawa M, Trifinopoulos J, Maurer B, Grausenburger R, Betram CA, Moriggl R, Rülicke T, Mullighan CG, Witalisz-Siepracka A, Walter W, Hoermann G, Sexl V, Gotthardt D (2024) A lineage-specific STAT5B^*N*642*H*^ mouse model to study NK-cell leukemia. Blood 143(24):2474–2489.

25. Suske T, Sorger H, Manhart G, Ruge F, Prutsch N, Zimmerman MW, Eder T, Abdallah DI, Maurer B, Wagner C, Schönefeldt S, Spirk K, Pichler A, Pemovska T, Schweicker C, Pölöske D, Hubanic E, Jungherz D, Müller TA, Aung MMK, Orlova A, Pham HTT, Zimmel K, Krausgruber T, Bock C, Müller M, Dahlhoff M, Boersma A, Rülicke T, Fleck R, de Araujo ED, Gunning PT, Aittokallio T, Mustjoki S, Sanda T, Hartmann S, Grebien F, Hoermann G, Haferlach T, Staber PB, Neubauer HA, Look AT, Herling M, Moriggl R (2024) Hyperactive STAT5 hijacks T cell receptor signaling and drives immature T cell acute lymphoblastic leukemia. Journal of Clinical Investigation 134(8):e168536.

26. Rajala HLM, Eldfors S, Kuusanmäki H, van Adrichem AJ, Olson T, Lagström S, Andersson EI, Jerez A, Clemente MJ, Yan Y, Zhang D, Awwad A, Ellonen P, Kallioniemi O, Wennerberg K, Porkka K, Maciejewski JP, Loughran TP, Heckman C, Mustjoki S (2013) Discovery of somatic STAT5b mutations in large granular lymphocytic leukemia. Blood 121(22):4541–4550.

27. Bandapalli OR, Schuessele S, Kunz JB, Rausch T, Stutz AM, Tal N, Geron I, Gershman N, Izraeli S, Eilers J, Vaezipour N, Kirschner-Schwabe R, Hof J, von Stackelberg A, Schrappe M, Stanulla M, Zimmermann M, Koehler R, Avigad S, Handgretinger R, Frismantas V, Bourquin JP, Bornhauser B, Korbel JO, Muckenthaler MU, Kulozik AE (2014) The activating STAT5B N642H mutation is a common abnormality in pediatric T-cell acute lymphoblastic leukemia and confers a higher risk of relapse. Haematologica 99(10):e188–e192.

28. Wingelhofer B, Maurer B, Heyes EC, Cumaraswamy AA, Berger-Becvar A, de Araujo ED, Orlova A, Freund P, Ruge F, Park J, Tin G, Ahmar S, Lardeau CH, Sadovnik I, Bajusz D, Keserű GM, Grebien F, Kubicek S, Valent P, Gunning PT, Moriggl R (2018) Pharmacologic inhibition of STAT5 in acute myeloid leukemia. Leukemia 32(5):1135–1146.

29. Cross NCP, Hoade Y, Tapper WJ, Carreno-Tarragona G, Fanelli T, Jawhar M, Naumann N, Pieniak I, Lübke J, Ali S, Bhuller K, Burgstaller S, Cargo C, Cavenagh J, Duncombe AS, Das-Gupta E, Evans P, Forsyth P, George P, Grimley C, Jack F, Munro L, Mehra V, Patel K, Rismani A, Sciuccati G, Thomas-Dewing R, Thornton P, Virchis A, Watt S, Wallis L, Whiteway A, Zegocki K, Bain BJ, Reiter A, Chase A (2018) Recurrent activating STAT5b N642H mutation in myeloid neoplasms with eosinophilia. Leukemia 33:415–425.

30. Ariyoshi K, Nosaka T, Yamada K, Onishi M, Oka Y, Miyajima A, Kitamura T (2000) Constitutive activation of STAT5 by a point mutation in the SH2 domain. Journal of Biological Chemistry 275(32):24407–24413.

31. Cordero-Morales JF, Jogini V, Lewis A, Vásquez V, Cortes DM, Roux B, Perozo E (2007) Molecular driving forces determining potassium channel slow inactivation. Nature Structural & Molecular Biology 14(11):1062–1069.

32. Hollingsworth SA, Dror RO (2018) Molecular dynamics simulation for all. Neuron 99(6):1129–1143.

33. Di Rita A, Angelini DF, Maiorino T, Caputo V, Cascella R, Kumar M, Tiberti M, Lambrughi M, Wesch N, Löhr F, Dötsch V, Carinci M, D’Acunzo P, Chiurchiù V, Papaleo E, Rogov VV, Giardina E, Battistini L, Strappazzon F (2021) Characterization of a natural variant of human NDP52 and its functional consequences on mitophagy. Cell Death & Differentiation 28(8):2499–2516.

34. Degn K, Beltrame L, Dahl Hede F, Sora V, Nicolaci V, Vabistsevits M, Schmiegelow K, Wadt K, Tiberti M, Lambrughi M, Papaleo E (2022) Cancer-related mutations with local or long-range effects on an allosteric loop of p53. Journal of Molecular Biology 434(17):167663.

35. Dölker N, Górna MW, Sutto L, Torralba AS, Superti-Furga G, Gervasio FL (2014) The SH2 domain regulates c-Abl kinase activation by a cyclin-like mechanism and remodulation of the hinge motion. PLoS Computational Biology 10(10):e1003863.

36. Anselmi M, Calligari P, Hub JS, Tartaglia M, Bocchinfuso G, Stella L (2020) Structural determinants of phosphopeptide binding to the N-terminal Src homology 2 domain of the SHP2 phosphatase. Journal of Chemical Information and Modeling 60(6):3157–3171.

37. Hobbs HT, Shah NH, Badroos JM, Gee CL, Marqusee S, Kuriyan J (2021) Differences in the dynamics of the tandem-SH2 modules of the Syk and ZAP-70 tyrosine kinases. Protein Science 30(12):2373–2384.

38. Marasco M, Kirkpatrick J, Carlomagno T, Hub JS, Anselmi M (2023) Experiment-guided molecular simulations define a heterogeneous structural ensemble for the PTPN11 tandem SH2 domains. Chemical Science 14:5743–5755.

39. Langenfeld F, Guarracino Y, Arock M, Trouvé A, Tchertanov L (2015) How intrinsic molecular dynamics control intramolecular communication in signal transducers and activators of transcription factor STAT5. PLoS ONE 10(12):e0145142.

40. Lin J, Buettner R, Yuan YC, Yip R, Horne D, Jove R, Vaidehi N (2009) Molecular dynamics simulations of the conformational changes in signal transducers and activators of transcription, Stat1 and Stat3. Journal of Molecular Graphics and Modelling 28(4):347–356.

41. Fahrenkamp D, Li J, Ernst S, de Leur HSV, Chatain N, Kuster A, Koschmieder S, Luscher B, Rossetti G, Muller-Newen G (2016) Intramolecular hydrophobic interactions are critical mediators of STAT5 dimerization. Scientific Reports 6:35454.

42. Gianti E, Zauhar RJ (2015) An SH2 domain model of STAT5 in complex with phospho-peptides define “STAT5 binding signatures”. Journal of Computer-Aided Molecular Design 29:451–470.

43. Ŕacz A, Mihalovits LM, Bajusz D, Héberger K, Miranda-Quintana RA (2022) Molecular dynamics simulations and diversity selection by extended continuous similarity indices. Journal of Chemical Information and Modeling 62(14):3415–3425.

44. Kaneko T, Huang H, Zhao B, Li L, Liu H, Voss CK, Wu C, Schiller MR, Li SSC (2010) Loops govern SH2 domain specificity by controlling access to binding pockets. Science Signaling 3(120):ra34.

45. Marasco M, Carlomagno T (2020) Specificity and regulation of phosphotyrosine signaling through SH2 domains. Journal of Structural Biology: X 4:100026.

46. Chen X, Vinkemeier U, Zhao Y, Jeruzalmi D, Darnell JE, Kuriyan J (1998) Crystal structure of a tyrosine phosphorylated STAT-1 dimer bound to DNA. Cell 93(5):827–839.

47. Huang Z, Liu H, Nix J, Xu R, Knoverek CR, Bowman GR, Amarasinghe GK, Sibley LD (2022) The intrinsically disordered protein TgIST from toxoplasma gondii inhibits STAT1 signaling by blocking cofactor recruitment. Nature Communications 13:4047.

48. Belo Y, Mielko Z, Nudelman H, Afek A, Ben-David O, Shahar A, Zarivach R, Gordan R, Arbely E (2019) Unexpected implications of STAT3 acetylation revealed by genetic encoding of acetyl-lysine. Biochimica et Biophysica Acta (BBA) -General Subjects 1863(9):1343–1350.

49. Liu BA, Engelmann BW, Nash PD (2012) The language of SH2 domain interactions defines phosphotyrosine-mediated signal transduction. FEBS Letters 586(17):2597–2605.

50. Bradshaw J, Mitaxov V, Waksman G (1999) Investigation of phosphotyrosine recognition by the SH2 domain of the Src kinase. Journal of Molecular Biology 293(4):971–985.

51. Waksman G, Kumaran S, Lubman O (2004) SH2 domains: role, structure and implications for molecular medicine. Expert Reviews in Molecular Medicine 6(3):1–18.

52. Singer AU (1998) Solution structure and electrostatic properties of an SH2 domain/phosphopeptide complex. PhD Thesis, University of Toronto.

53. Liu BA, Jablonowski K, Raina M, Arće M, Pawson T, Nash PD (2006) The human and mouse complement of SH2 domain proteins—establishing the boundaries of phosphotyrosine signaling. Molecular Cell 22(6):851–868.

54. Eck MJ, Atwell SK, Shoelson SE, Harrison SC (1994) Structure of the regulatory domains of the Src-family tyrosine kinase Lck. Nature 368(6473):764–769.

55. Taylor JD, Ababou A, Fawaz RR, Hobbs CJ, Williams MA, Ladbury JE (2008) Structure, dynamics, and binding thermodynamics of the v-Src SH2 domain: Implications for drug design. Proteins: Structure, Function, and Bioinformatics 73(4):929–940.

56. Singer AU, Forman-Kay JD (1997) pH titration studies of an SH2 domain-phosphopeptide complex: Unusual histidine and phosphate pKa values. Protein Science 6(9):1910–1919.

57. Rauscher S, Pomès R (2010) Molecular simulations of protein disorder. Biochemistry and Cell Biology 88(2):269–290.

58. Liao SM, Du QS, Meng JZ, Pang ZW, Huang RB (2013) The multiple roles of histidine in protein interactions. Chemistry Central Journal 7:44.

59. Guljas A (2021) Energetics of *π − π* stacking interactions: Implications in the phase separation of intrinsically disordered proteins. MSc Thesis, University of Toronto.

60. Yang Z, Wang Z, Tian X, Xiu P, Zhou R (2012) Amino acid analogues bind to carbon nanotube via *π*-*π* interactions: Comparison of molecular mechanical and quantum mechanical calculations. The Journal of Chemical Physics 136(2):025103.

61. Machida K, Mayer BJ (2005) The SH2 domain: versatile signaling module and pharmaceutical target. Biochimica et Biophysica Acta (BBA) -Proteins and Proteomics 1747:1–25.

62. Mao X, Ren Z, Parker GN, Sondermann H, Pastorello MA, Wang W, McMurray JS, Demeler B, Darnell JE, Chen X (2005) Structural bases of unphosphorylated STAT1 association and receptor binding. Molecular Cell 17(6):761–771.

63. Neculai D, Neculai AM, Verrier S, Straub K, Klumpp K, Pfitzner E, Becker S (2005) Structure of the unphosphorylated STAT5a dimer. Journal of Biological Chemistry 280(49):40782–40787.

64. Wilson DJ, Konermann L (2003) A capillary mixer with adjustable reaction chamber volume for millisecond time-resolved studies by electrospray mass spectrometry. Analytical Chemistry 75(23):6408–6414.

65. Lento C, Zhu S, Brown KA, Knox R, Liuni P, Wilson DJ (2017) Time-resolved ElectroSpray ionization hydrogen-deuterium exchange mass spectrometry for studying protein structure and dynamics. Journal of Visualized Experiments 122:e55464.

66. Bhattacharya D, Teramo A, Gasparini VR, Huuhtanen J, Kim D, Theodoropoulos J, Schiavoni G, Barilà G, Vicenzetto C, Calabretto G, Facco M, Kawakami T, Nakazawa H, Falini B, Tiacci E, Ishida F, Semenzato G, Kelkka T, Zambello R, Mustjoki S (2022) Identification of novel STAT5B mutations and characterization of TCR*β* signatures in CD4+ T-cell large granular lymphocyte leukemia. Blood Cancer Journal 12:31.

67. Huang J, Rauscher S, Nawrocki G, Ran T, Feig M, de Groot BL, Grubmüller H, MacKerell AD (2017) CHARMM36m: an improved force field for folded and intrinsically disordered proteins. Nature Methods 14:71–73.

68. MacKerell AD, Bashford D, Bellott M, Dunbrack RL, Evanseck JD, Field MJ, Fischer S, Gao J, Guo H, Ha S, Joseph-McCarthy D, Kuchnir L, Kuczera K, Lau FTK, Mattos C, Michnick S, Ngo T, Nguyen DT, Prodhom B, Reiher WE, Roux B, Schlenkrich M, Smith JC, Stote R, Straub J, Watanabe M, Wíorkiewicz-Kuczera J, Yin D, Karplus M (1998) All-atom empirical potential for molecular modeling and dynamics studies of proteins. The Journal of Physical Chemistry B 102(18):3586–3616.

69. Hess B, Bekker H, Berendsen HJC, Fraaije JGEM (1997) LINCS: A linear constraint solver for molecular simulations. Journal of Computational Chemistry 18(12):1463–1472.

70. Miyamoto S, Kollman PA (1992) Settle: An analytical version of the SHAKE and RATTLE algorithm for rigid water models. Journal of Computational Chemistry 13(8):952–962.

71. Parrinello M, Rahman A (1981) Polymorphic transitions in single crystals: A new molecular dynamics method. Journal of Applied Physics 52(12):7182–7190.

72. Bussi G, Donadio D, Parrinello M (2007) Canonical sampling through velocity rescaling. The Journal of Chemical Physics 126(1):014101.

73. Waterhouse A, Bertoni M, Bienert S, Studer G, Tauriello G, Gumienny R, Heer FT, de Beer TAP, Rempfer C, Bordoli L, Lepore R, Schwede T (2018) SWISS-MODEL: homology modelling of protein structures and complexes. Nucleic Acids Research 46:W296–W303.

74. Chen VB, Arendall WB, Headd JJ, Keedy DA, Immormino RM, Kapral GJ, Murray LW, Richardson JS, Richardson DC (2009) MolProbity: all-atom structure validation for macromolecular crystallography. Acta Crystallographica Section D 66:12–21.

75. Fiser A, Do RKG, S^̌^ali A (2000) Modeling of loops in protein structures. Protein Science 9(9):1753– 1773.

76. Pettersen EF, Goddard TD, Huang CC, Couch GS, Greenblatt DM, Meng EC, Ferrin TE (2004) UCSF Chimera -a visualization system for exploratory research and analysis. Journal of Computational Chemistry 25(13):1605–1612.

77. Jo S, Kim T, Iyer VG, Im W (2008) CHARMM-GUI: A web-based graphical user interface for CHARMM. Journal of Computational Chemistry 29(11):1859–1865.

78. Jo S, Cheng X, Islam SM, Huang L, Rui H, Zhu A, Lee HS, Qi Y, Han W, Vanommeslaeghe K, MacKerell AD, Roux B, Im W (2014) CHARMM-GUI PDB manipulator for advanced modeling and simulations of proteins containing nonstandard residues. Advances in Protein Chemistry and Structural Biology 96:235–265.

79. Berendsen H, van der Spoel D, van Drunen R (1995) GROMACS: A message-passing parallel molecular dynamics implementation. Computer Physics Communications 91(1-3):43–56.

80. Berendsen HJC, Postma JPM, van Gunsteren WF, DiNola A, Haak JR (1984) Molecular dynamics with coupling to an external bath. The Journal of Chemical Physics 81(8):3684–3690.

81 Schrödinger, LLC. The PyMOL molecular graphics system, version 2.3.

82. Søndergaard CR, Olsson MHM, Rostkowski M, Jensen JH (2011) Improved treatment of ligands and coupling effects in empirical calculation and rationalization of pK*_a_* values. Journal of Chemical Theory and Computation 7:2284–2295.

83. Darden T, York D, Pedersen L (1993) Particle mesh Ewald: An N*·*log(N) method for Ewald sums in large systems. The Journal of Chemical Physics 98(12):10089–10092.

84. Guex N, Peitsch MC (1997) SWISS-MODEL and the Swiss-PdbViewer: An environment for comparative protein modeling. Electrophoresis 18(15):2714–2723.

85. de Groot B, Hayward S, van Aalten D, Amadei A, Berendsen H (1998) Domain motions in bacteriophage T4 lysozyme: A comparison between molecular dynamics and crystallographic data. Proteins: Structure, Function, and Genetics 31(2):116–127.

86. Peters JH, de Groot BL (2012) Ubiquitin dynamics in complexes reveal molecular recognition mechanisms beyond induced fit and conformational selection. PLoS Computational Biology 8(10):e1002704.

87. Buelens FP, Leonov H, de Groot BL, Grubmüller H (2021) ATP–magnesium coordination: Protein structure-based force field evaluation and corrections. Journal of Chemical Theory and Computation 17(3):1922–1930.

88. Konagurthu AS, Whisstock JC, Stuckey PJ, Lesk AM (2006) MUSTANG: A multiple structural alignment algorithm. Proteins 64(3):559–574.

89. Pedregosa F, Varoquaux G, Gramfort A, Michel V, Thirion B, Grisel O, Blondel M, Prettenhofer P, Weiss R, Dubourg V, Vanderplas J, Passos A, Cournapeau D, Brucher M, Perrot M, Duchesnay E. Scikit-learn: Machine learning in Python. Journal of Machine Learning Research 12:2825–2830.

90. Humphrey W, Dalke A, Schulten K (1996) VMD: Visual molecular dynamics. Journal of Molecular Graphics 14(1):33–38.

91. Kabsch W, Sander C (1983) Dictionary of protein secondary structure: Pattern recognition of hydrogen-bonded and geometrical features. Biopolymers 22(12):2577–2637.

92. McGibbon RT, Beauchamp KA, Harrigan MP, Klein C, Swails JM, Herńandez CX, Schwantes CR, Wang LP, Lane TJ, Pande VS (2015) MDtraj: A modern open library for the analysis of molecular dynamics trajectories. Biophysical Journal 109(8):1528 – 1532.

93. Shrake A, Rupley J (1973) Environment and exposure to solvent of protein atoms. lysozyme and insulin. Journal of Molecular Biology 79(2):351–371.

94. Michaud-Agrawal N, Denning EJ, Woolf TB, Beckstein O (2011) MDAnalysis: A toolkit for the analysis of molecular dynamics simulations. Journal of Computational Chemistry 32(10):2319–2327.

95. Zhou QY, Park J, Koltun V (2018) Open3D: A modern library for 3D data processing. arXiv.

96. Ball DP, Lewis AM, Williams D, Resetca D, Wilson DJ, Gunning PT (2016) Signal transducer and activator of transcription 3 (STAT3) inhibitor, S3I-201, acts as a potent and non-selective alkylating agent. Oncotarget 7(15):20669–20679.

97. Knox R, Lento C, Wilson DJ (2018) Mapping conformational dynamics to individual steps in the TEM-1 *β*-lactamase catalytic mechanism. Journal of Molecular Biology 430:3311–3322.

98. Rey M, Sarpe V, Burns KM, Buse J, Baker CA, van Dijk M, Wordeman L, Bonvin AM, Schriemer DC (2014) Mass spec studio for integrative structural biology. Structure 22(10):1538–1548.

