## Supporting Information for "The structural influence of the oncogenic driver mutation N642H in the STAT5B SH2 domain"

### Contents

### List of Figures

S1 Domain organization of STAT5B. . . . . . . . . . . . . . . . . . . . . . . . . . . . . . . S3
S2 Predicted disorder in human STAT proteins. . . . . . . . . . . . . . . . . . . . . . . . . S4
S3 Conformational landscape of mammalian STAT SH2 domains. . . . . . . . . . . . . . . S5
S4 STAT SH2 conformational landscape - principal components . . . . . . . . . . . . . . . S6
S5 Contacts with residue pY699 in the parallel dimer . . . . . . . . . . . . . . . . . . . . S7
S6 Contacts stabilizing the parallel dimer interface. . . . . . . . . . . . . . . . . . . . . . S8
S7 Binding pocket changes in the parallel dimer . . . . . . . . . . . . . . . . . . . . . . . S9
S8 Structural differences between STAT5B and STAT5B$^{N642H}$ SH2 domains . . . . . . S10
S9 Structural differences between the STAT5B and STAT5B$^{N642H}$ parallel dimers in the phosphotyrosine motif region. . . . . . . . . . . . . . . . . . . . . . . . . . . . . . . S11
S10 Relative orientations of N642H and pY699. . . . . . . . . . . . . . . . . . . . . . . . . S12
S11 R618 and N642(H) hydrogen bonds in the apo systems. . . . . . . . . . . . . . . . . S13
S12 B-factors from STAT5B crystal structures. . . . . . . . . . . . . . . . . . . . . . . . . S14
S13 STAT5B SH2 structural ensembles. . . . . . . . . . . . . . . . . . . . . . . . . . . . . S15
S14 Apo STAT5B ensembles projected onto the PC space of the parallel dimer and STAT5B$^{N642H}$ antiparallel dimer trajectories. . . . . . . . . . . . . . . . . . . . . . . S16
S15 STAT1 SH2 domain has two $\beta$-sheet states. . . . . . . . . . . . . . . . . . . . . . . . S17
S16 STAT5A SH2 domain has two $\beta$-sheet states. . . . . . . . . . . . . . . . . . . . . . . S18
S17 Sequence alignment of STAT SH2 domains. . . . . . . . . . . . . . . . . . . . . . . . . S19

### List of Tables

S1 List of simulations . . . . . . . . . . . . . . . . . . . . . . . . . . . . . . . . . . . . . S20
S2 List of crystal structures of mammalian STAT SH2 domains . . . . . . . . . . . . . . S21

**a**

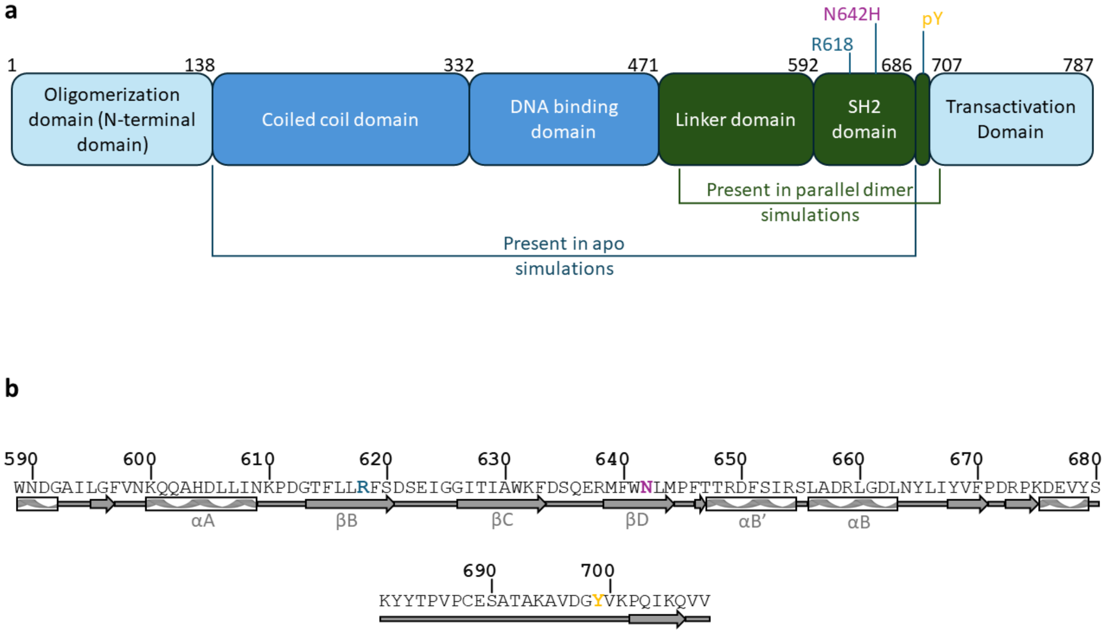


**Figure S1: Domain organization of STAT5B.** (a) A schematic showing the domain organization of STAT5B. The domain boundaries are as defined in de Araujo *et al.*[1]. Key residues R618, N642(H), and pY699 are highlighted. Residues present in the simulations of the parallel dimer and apo systems are indicated. The phosphotyrosine motif region (residues 687-706) is found between the SH2 domain and the transactivation domain. (b) Part of the sequence of STAT5B is shown (UniProt ID P51692, residues 589-708). This sequence includes the SH2 domain and the phosphotyorsine motif region. The secondary structure is shown below the sequence and residue numbers are shown above. Important residues are coloured as follows: R618 – blue, N642 – magenta, and Y699 – yellow.

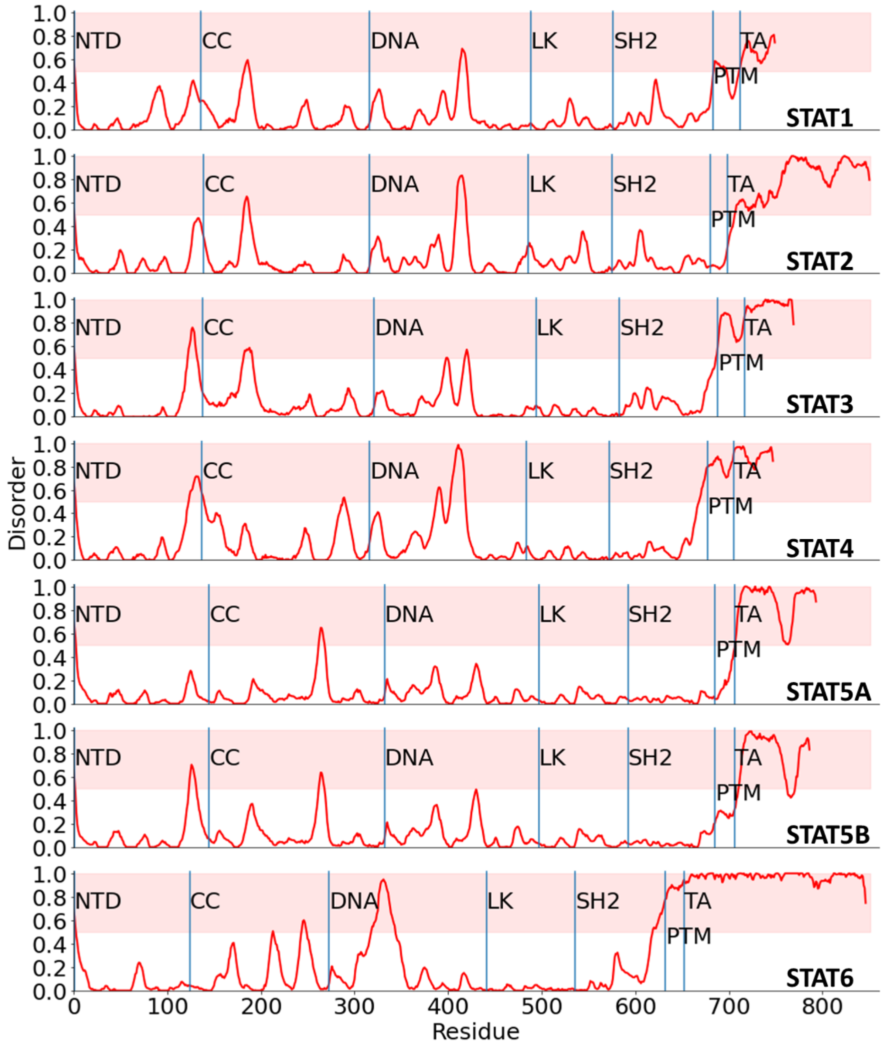


**Figure S2: Predicted disorder in human STAT proteins.** The predicted disorder by residue of each human STAT protein, as predicted by Metapredict V2[2;3]. Higher values indicate a higher likelihood of the residue being disordered. Above a threshold value of 0.5, the residue is predicted to be disordered[2;3], indicated by the red shading. Domain boundaries of each protein as defined by Lim *et al.*[4] are indicated. NTD - N-terminal Domain, CC - coiled coil domain, DNA - DNA binding domain, LK - linker domain, SH2 - Src-homology 2 domain, PTM - phosphotyrosine motif region, TA - transactivation domain.

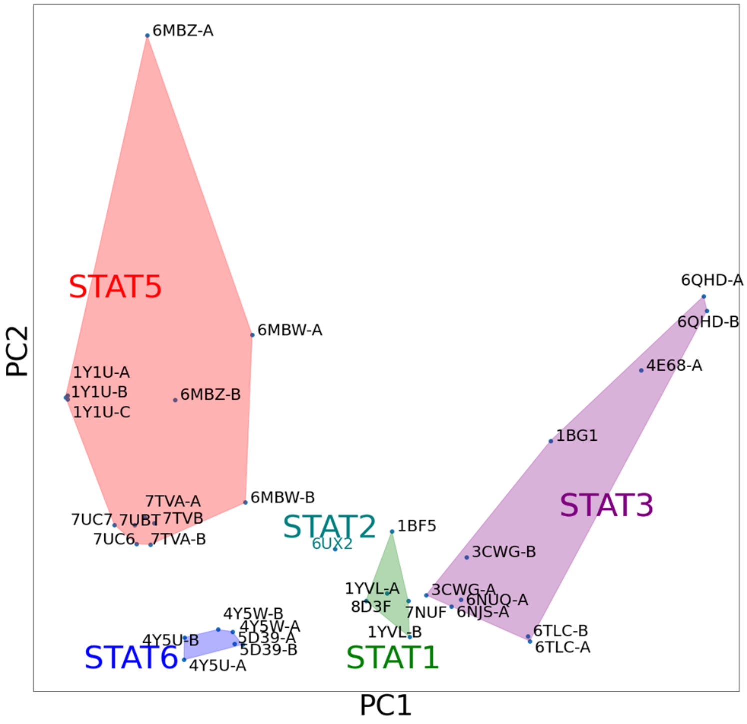


**Figure S3: Conformational landscape of mammalian STAT SH2 domains.** PCA of all available crystal structures of mammalian STAT SH2 domains. The pairwise distance between C$\alpha$ atoms was used as a feature vector (see Methods). Each structure is projected onto the 2D space defined by the first two principal components, PC1 and PC2. Structures are labelled with the corresponding PDB ID. For cases where there are multiple chains in the crystal structure, they are labelled A, B, and C. Shaded regions indicate where different STAT proteins reside in the conformational space.

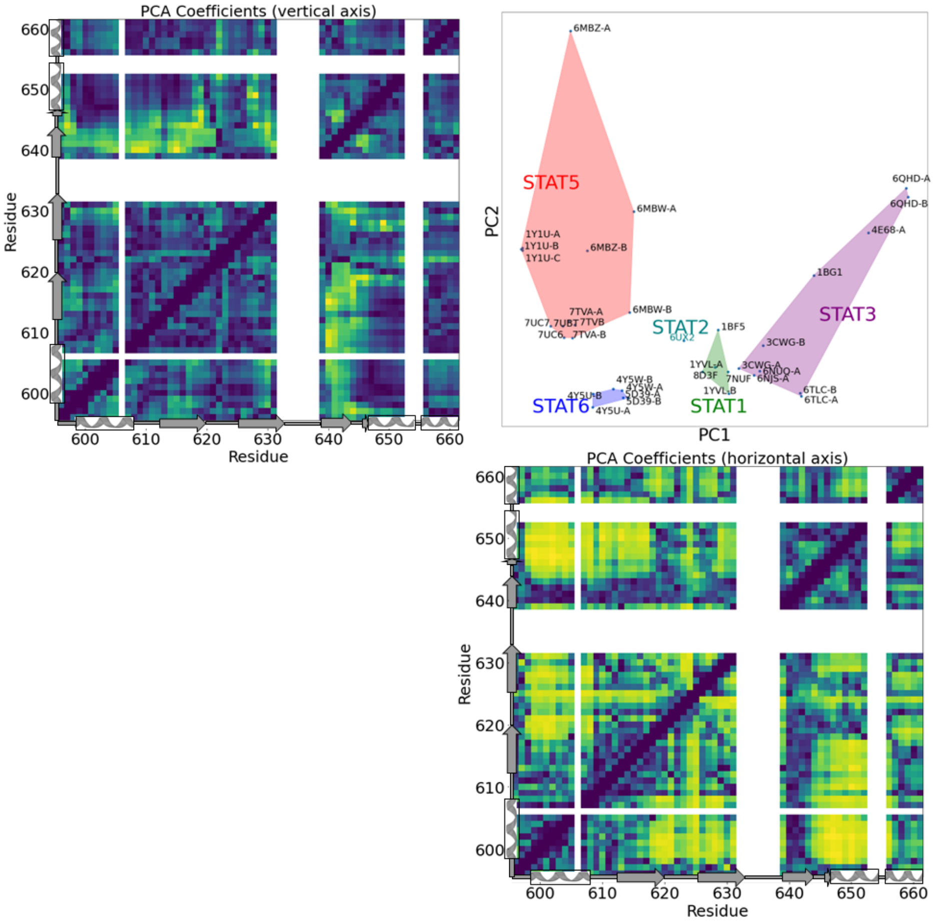


**Figure S4: STAT SH2 conformational landscape - principal components.** The contributions to the first two principal components in the PCA shown in Figure S3. PC1 and PC2 account for 35% and 10% of the total variance in the data, respectively. The contributions to PC1 are shown below the PCA space and the contributions to PC2 are shown to the left of the PCA space. Because pairwise distances between C$\alpha$ atoms were used as the feature in PCA, the contributions of each of these pairwise distances is shown, with yellow indicating a larger contribution, and dark blue indicating a smaller contribution. Green indicates intermediate contribution. Regions in white indicate residues that did not align between all STAT SH2 domain crystal structures and were, therefore, not included in the PCA.

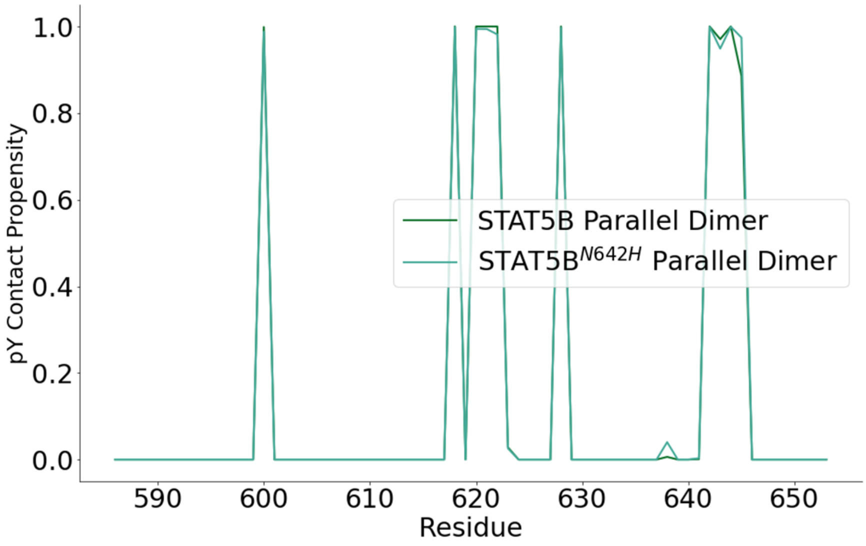


**Figure S5: Contacts with residue pY699 in the parallel dimer.** The portion of simulation frames that each residue in the SH2 domain is in contact with residue pY699 of the opposite monomer. Contact frequencies are provided for both the wild-type and N642H parallel dimer simulations. Residues in the SH2 domain with the most probable interactions with pY699 include K600, R618, S620, D621, S622, T628, N642/H642, L643, M644, and P645. These residues comprise the pY pocket. Note that the pY contact propensities are very similar between STAT5B and STAT5B$^{N642H}$, so the lines overlap for most residues.

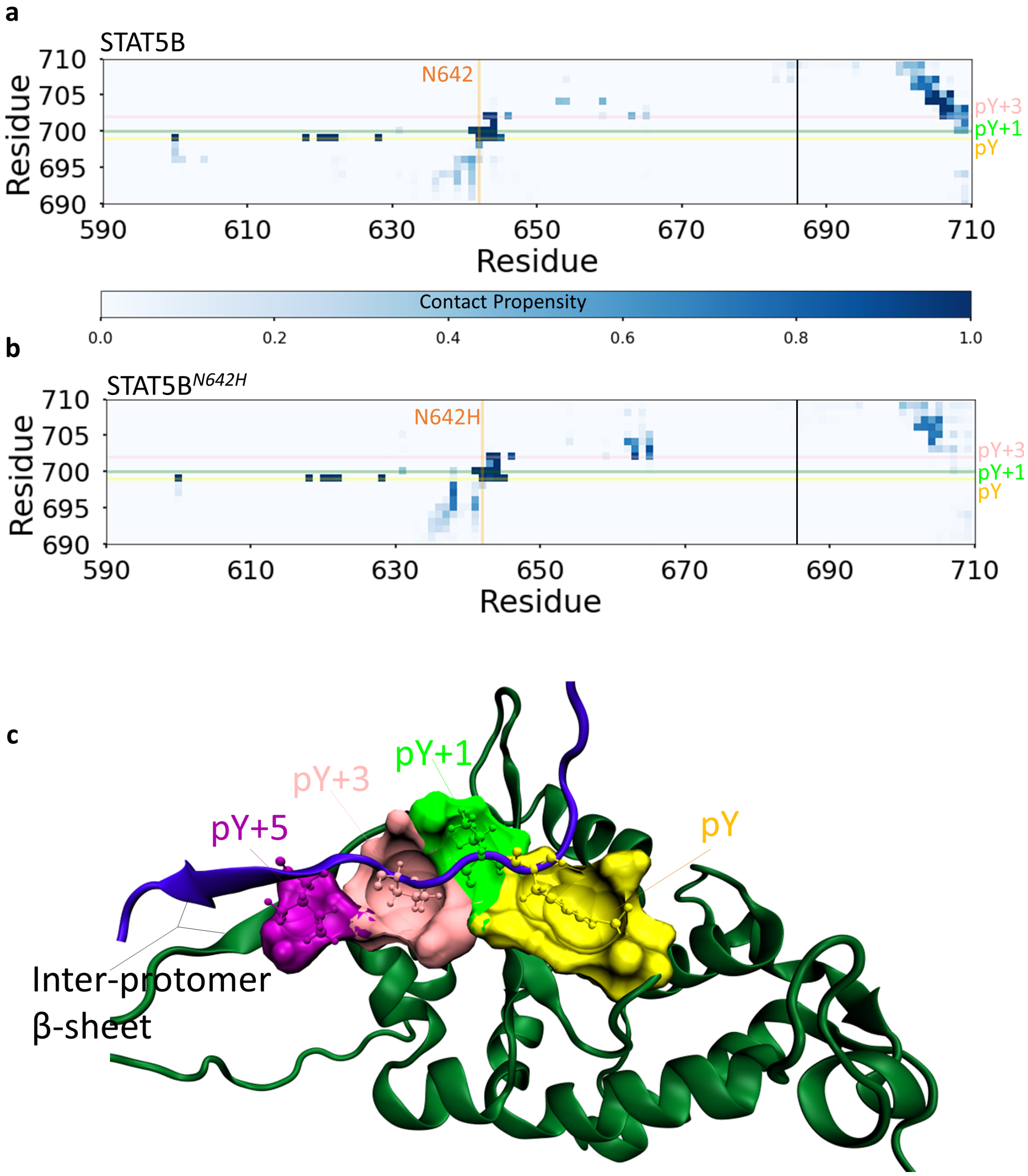


**Figure S6: Contacts stabilizing the parallel dimer interface.** Pairwise residue contact propensities in the wild type (a) and STAT5B$^{N642H}$ (b) parallel dimer simulations. Residue numbers on the horizontal axis correspond to the SH2 domain and phosphotyrosine motif region (separated by a black vertical line) of one monomer in the dimer, while residue numbers on the vertical axis correspond to the phosphotyrosine motif region of the opposite monomer. In panels (a) and (b), residue N642(H) is indicated by a vertical orange line. pY, pY+1, and pY+3 residues are indicated by yellow, green, and pink horizontal lines, respectively. The colour of each square indicates the frequency of contact formation, with darker blue indicating higher propensity contacts. (c) A representative 3D structure of the parallel dimer interface. The SH2 domain is shown in green cartoon representation. pY, pY+1, pY+3 and pY+5 are shown in ball and stick representation, with residues from the opposite protomer within 3.5 Å of these residues shown in surface representation.

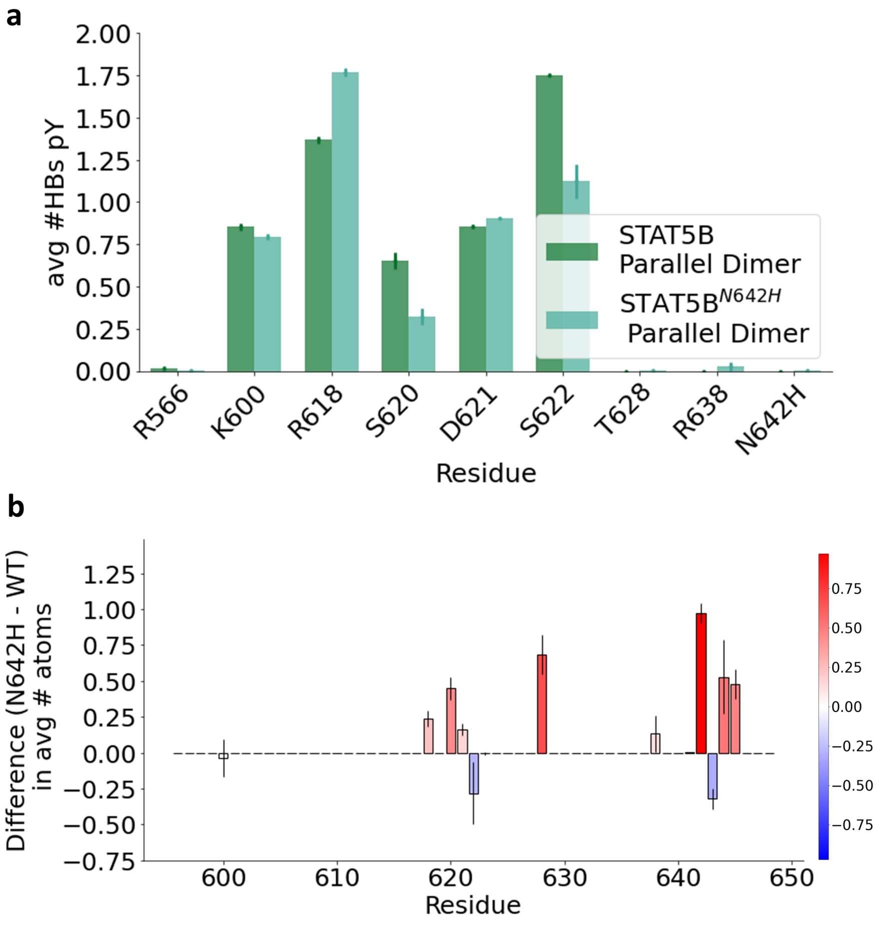


**Figure S7: Binding pocket changes in the parallel dimer** (a) Average number of hydrogen bonds between residue pY699 and other SH2 domain residues in simulations of STAT5B (dark green) and STAT5B$^{N642H}$ (light green) parallel dimers. (b) Average difference (mutant - wild type) in number of atoms in contact with pY. Red bars indicate more atoms in contact in the N642H parallel dimer, blue bars indicate more atoms in contact in the wild-type parallel dimer. Error bars indicate standard error of the mean obtained by treating each chain as an independent measurement.

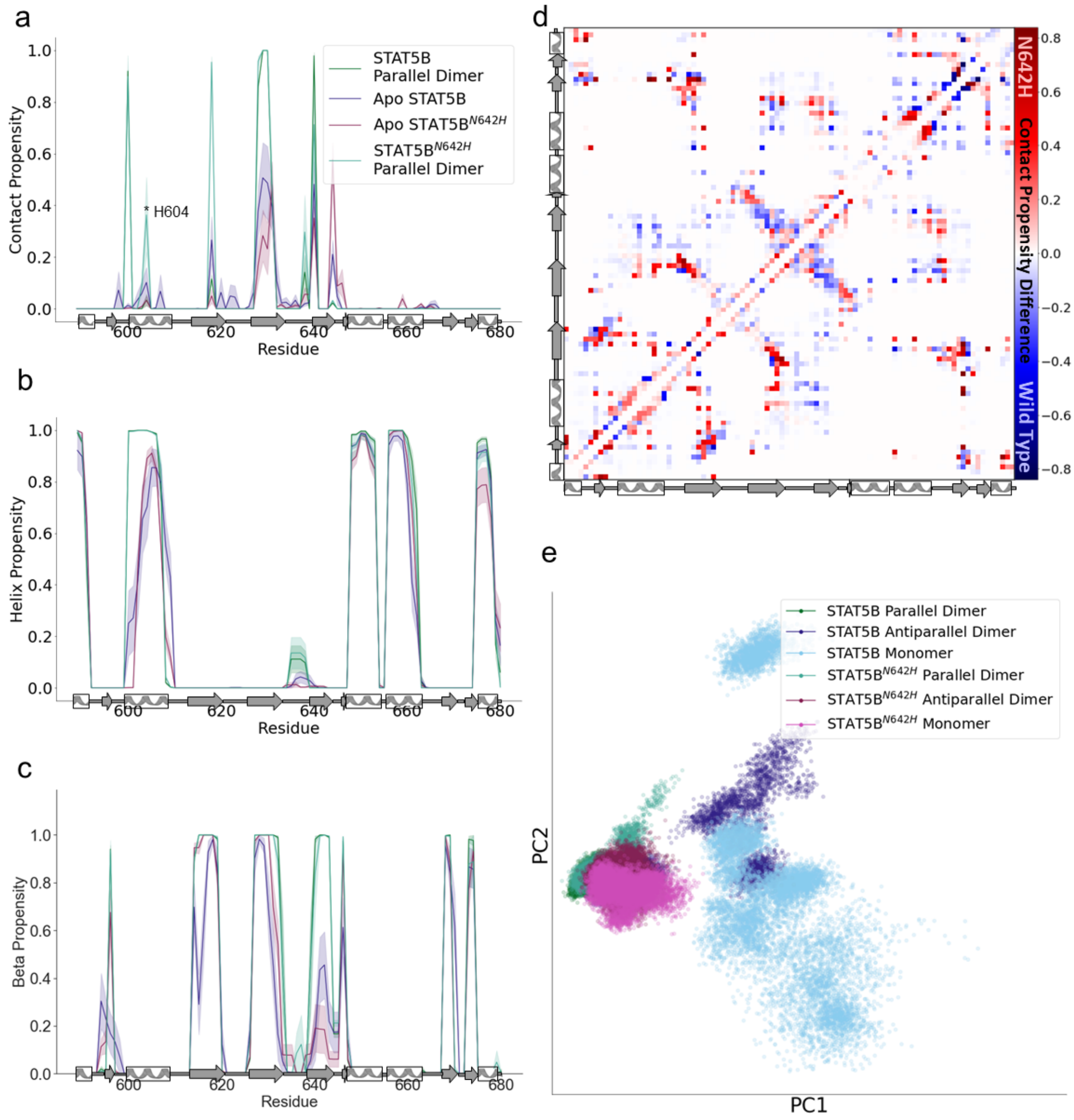


**Figure S8: Structural differences between STAT5B and STAT5B$^{N642H}$ SH2 domains.** (a) Contact propensity between the mutation site residue 642 and all other SH2 domain residues. The contact propensity with residue H604 is 0.034 $\pm$ 0.022 in the STAT5B parallel dimer and 0.363 $\pm$ 0.145 in the STAT5B$^{N642H}$ parallel dimer. The legend in this panel applies to panels (a) to (c). (b) Average helix propensity per residue in simulation trajectories. (c) Average beta propensity per residue in simulation trajectories. In (a)-(c), shaded regions indicate standard error of the mean obtained by treating each SH2 domain in the simulations as an independent measurement. (d) Average contact differences between apo STAT5B and STAT5B$^{N642H}$ SH2 domains. Red indicates that a contact is formed more often in the N642H mutant and blue indicates that a contact is formed more often in wild type. (e) Principal component analysis on pairwise C$\alpha$ distances for the SH2 domain for every frame of each simulation trajectory.

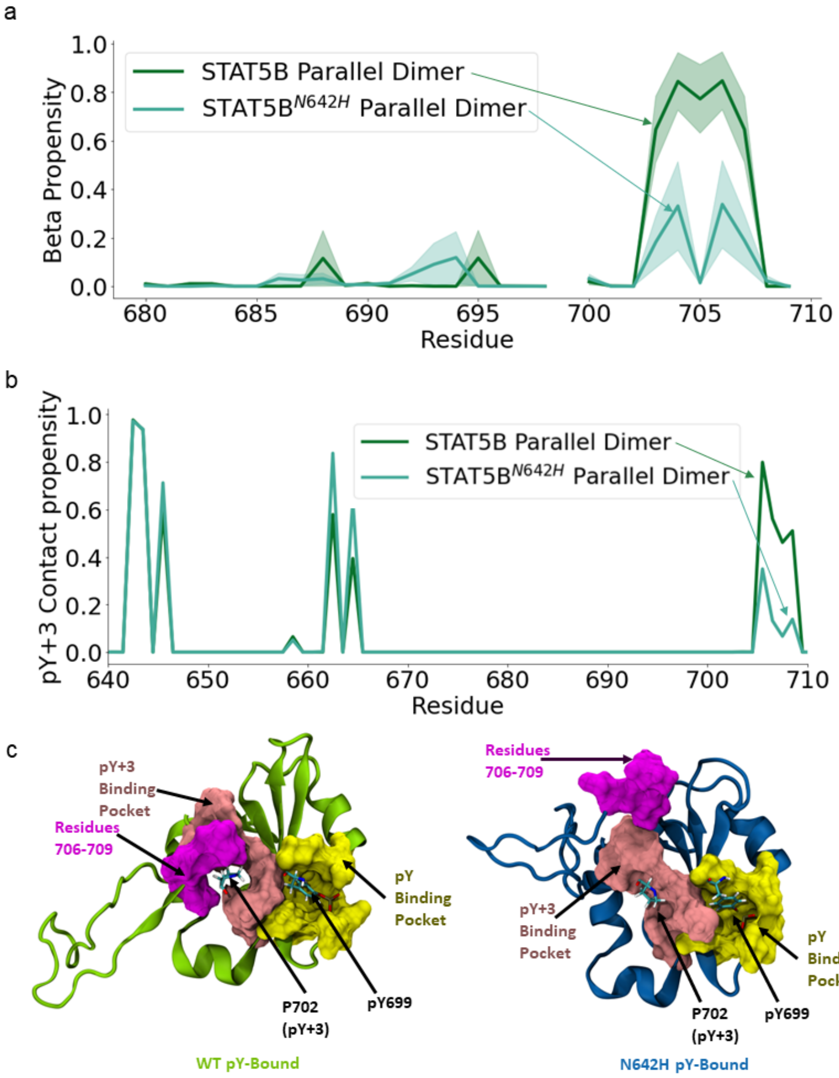


**Figure S9: Structural differences between the STAT5B and STAT5B$^{N642H}$ parallel dimers in the phosphotyrosine motif region.** (a) Beta propensity in the phosphotyrosine motif region (residues 680-708) in the simulations of the STAT5B and STAT5B$^{N642H}$ parallel dimers. Shaded regions indicate standard error of the mean obtained by treating each SH2 domain in the simulations as an independent measurement. (b) pY+3 contact propensity, which is the portion of simulation frames that each residue in the SH2 domain is in contact with the pY+3 residue (P702) of the opposite monomer in simulations of the parallel dimer. Results for STAT5B and STAT5B$^{N642H}$ parallel dimer simulations are indicated in dark green and light green, respectively. Residues in the SH2 domain with the most probable interactions with pY+3 include L643, M644, F646, L663, and Y665. Residues 706-709 in the phosphotyrosine motif region also interact with pY+3. (c) 3D structures showing representative conformations of the SH2 domain and phosphotyrosine motif region in the parallel dimer for STAT5B (left) and STAT5B$^{N642H}$ (right). The pY binding pocket (yellow) and pY+3 binding pocket (pink) residues are shown in surface representation. Residues 706-709 are shown in magenta. The pY and pY+3 residues from the opposite monomer are shown in licorice representation.

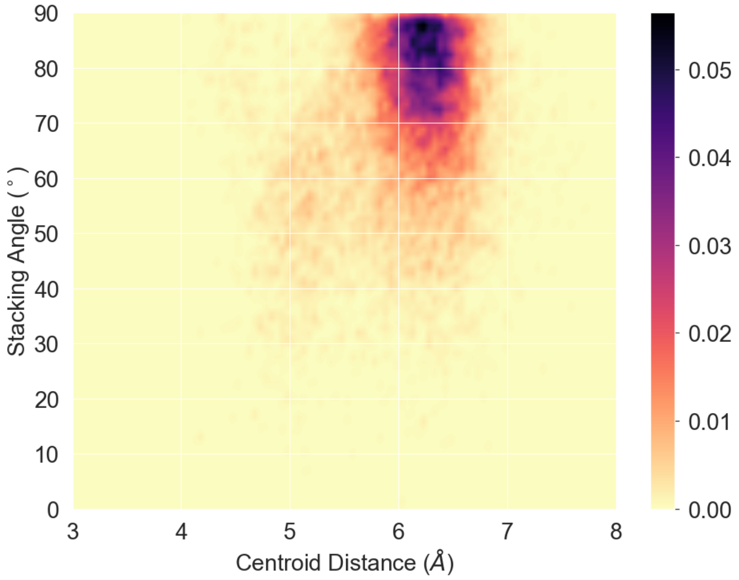


**Figure S10: Relative orientations of N642H and pY699.** The joint distribution of ring centroid distance and stacking angle is shown. The horizontal axis shows the centroid distance between the ring of pY699 and N642H. The vertical axis shows the stacking angle between the two rings, defined as the angle between the normal vectors to the planes defined by each ring. Distances and angles are computed using the STAT5B$^{N642H}$ parallel dimer simulations.

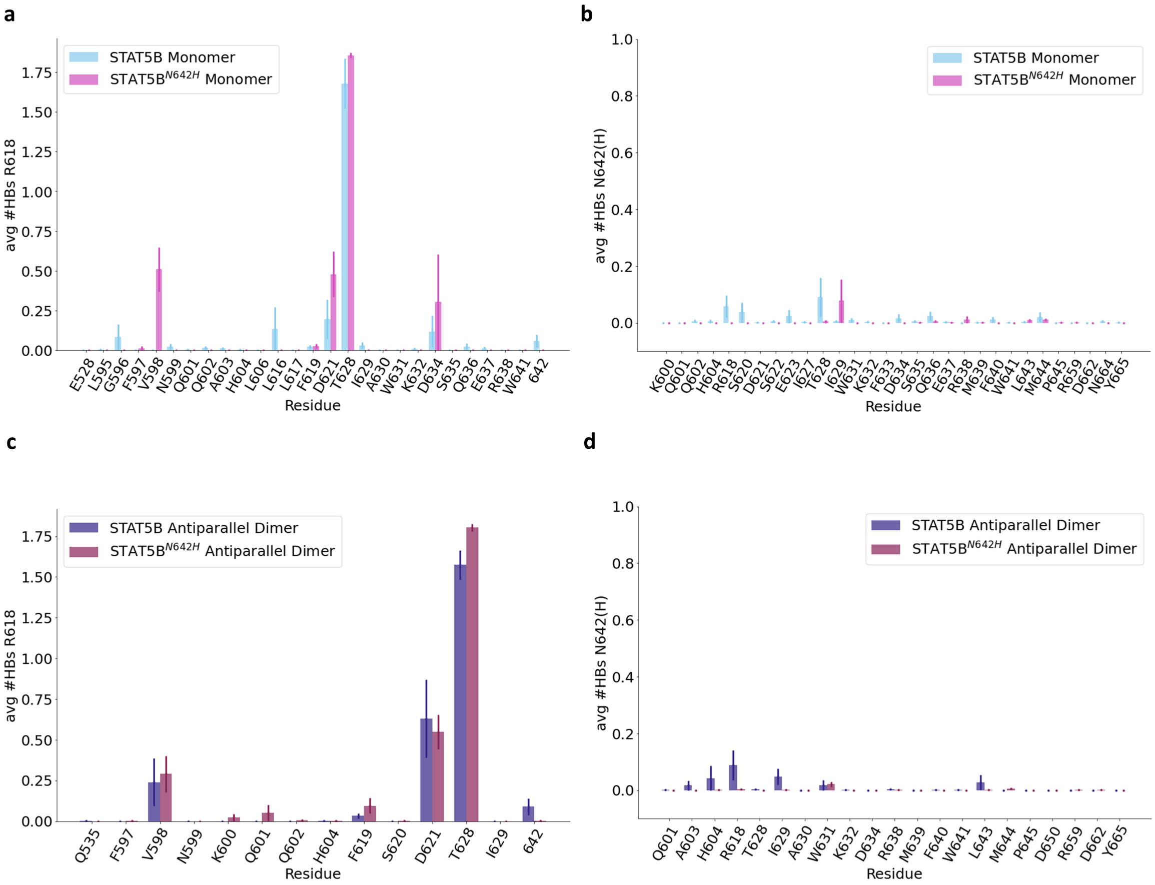


**Figure S11: R618 and N642(H) hydrogen bonds in the apo systems.** The probability of a hydrogen bond between R618 (a, c) and N642(H) (b, d) and other SH2 domain residues. Results are shown for simulations of STAT5B and STAT5B$^{N642H}$ monomers (a, b) and antiparallel dimers (c, d). In all figure panels, only residues with a non-zero probability of hydrogen bond formation with R618 or N642(H) are included. Vertical lines indicate the standard error of the mean treating each chain in each simulation as independent.

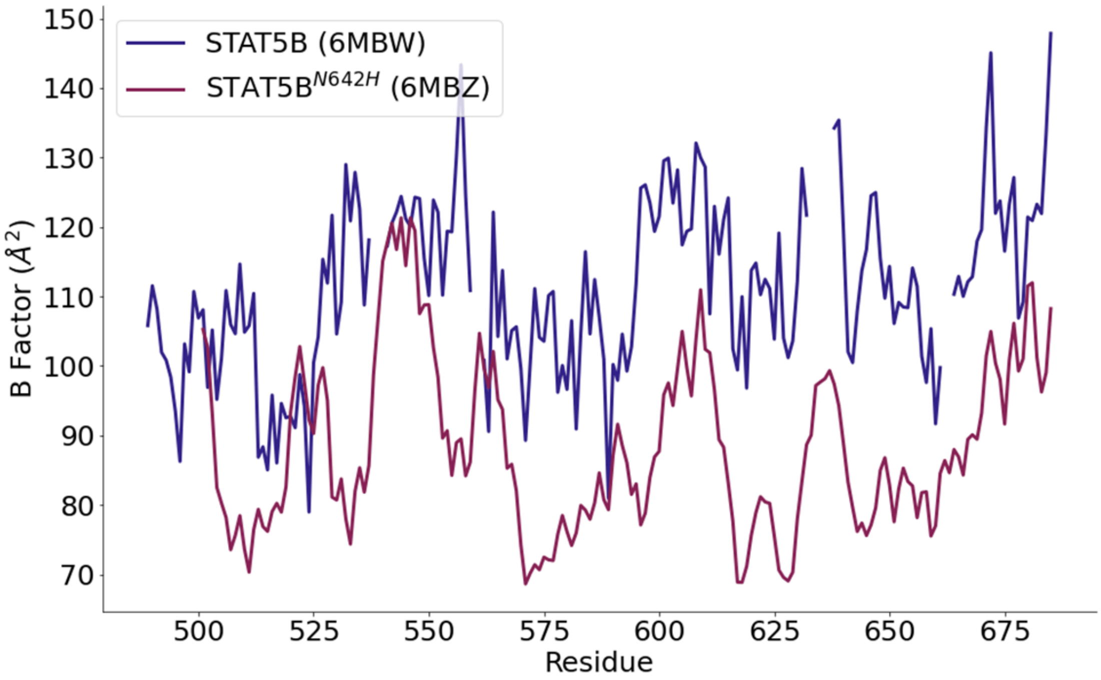


**Figure S12: B-factors from STAT5B crystal structures** B-factors from the crystal structures of STAT5B and STAT5B$^{N642H}$ antiparallel dimers[1] (PDB structures 6MBW and 6MBZ, which are crystal structures of the wild type and N642H mutant). Only the residues of the linker and SH2 domains are shown.

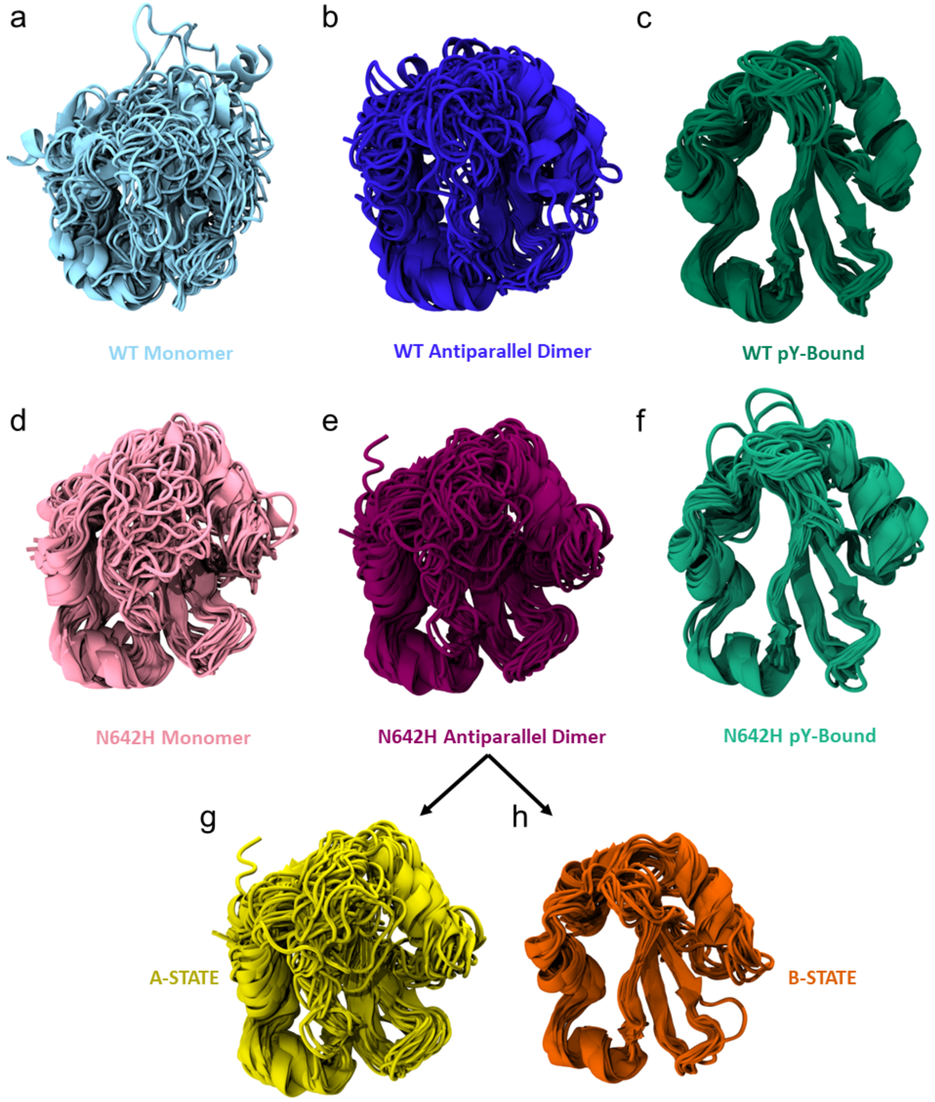


**Figure S13: STAT5B SH2 structural ensembles.** Ensemble of 24 structures randomly sampled from the MD simulation trajectories of each system, (a) wild-type pY-bound parallel dimer, (b) wild-type antiparallel dimer, (c) wild-type monomer, (d) N642H mutant pY-bound parallel dimer, (e) N642H mutant antiparallel dimer, and (f) N642H mutant monomer. The N642H mutant antiparallel dimer is also shown with ensembles corresponding to the A-state (g) and B-state (h).

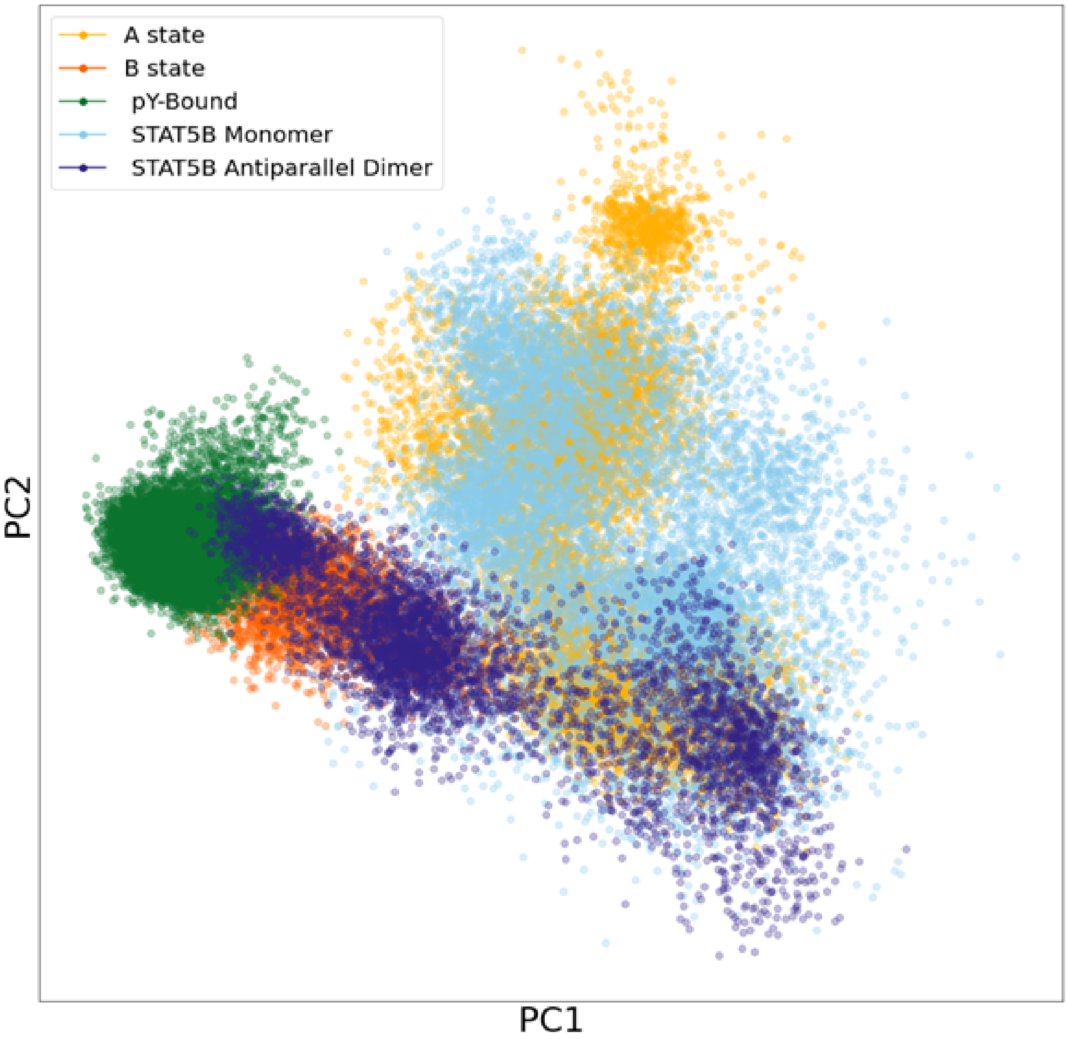


**Figure S14: Apo STAT5B ensembles projected onto the PC space of the parallel dimer and STAT5B$^{N642H}$ antiparallel dimer trajectories.** Every structure from the apo STAT5B simulation trajectories was projected onto the conformational space of the parallel dimer and STAT5B$^{N642H}$ antiparallel dimer SH2 domains. This is the same PC space as shown in Figure 4d. Each point represents one structure. The structures from each ensemble are coloured as indicated in the legend.

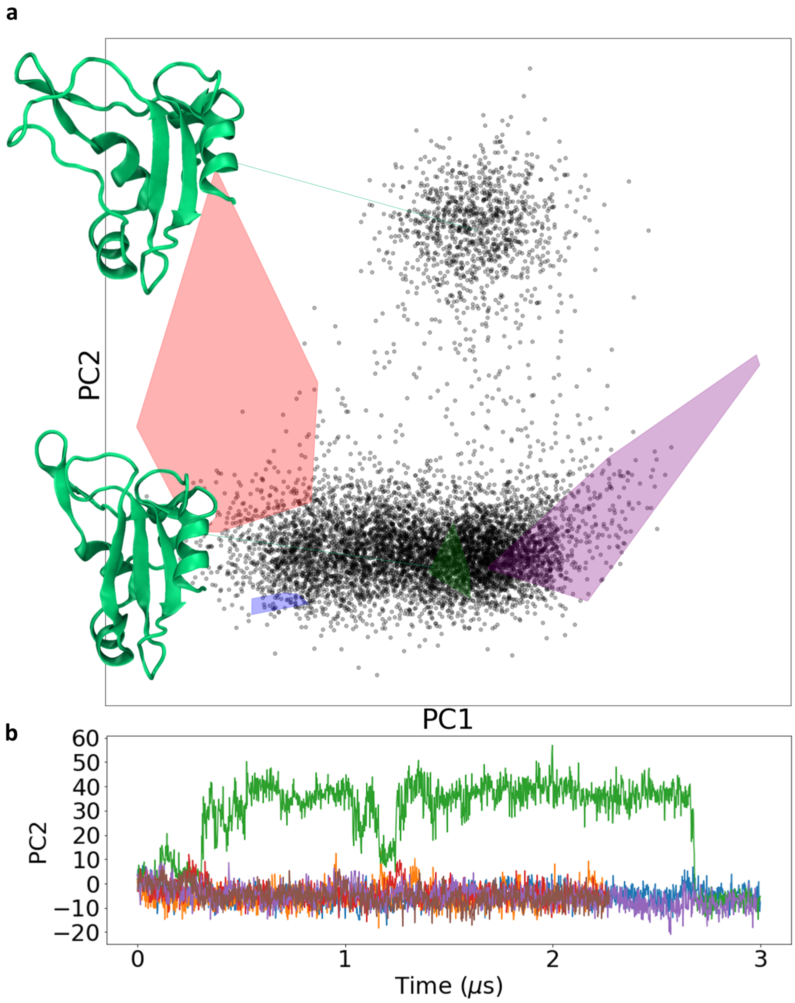


**Figure S15: STAT1 SH2 domain has two $\beta$-sheet states.** (a) STAT1 trajectories projected on the STAT SH2 conformational landscape. Representative structures from each state are provided (upper: $\beta$D strand in an open state, lower: $\beta$D strand forming contacts in a $\beta$-sheet). (b) Time series of PC2 for each STAT1 trajectory is shown. Each of the six trajectories is indicated in a different colour.

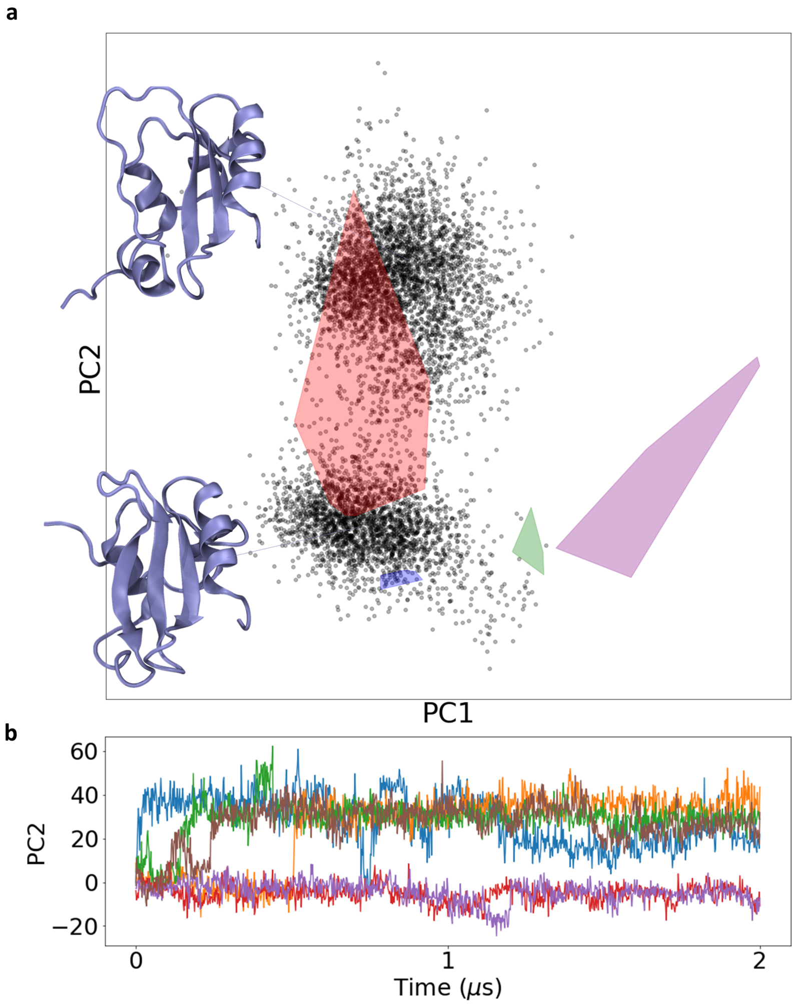


**Figure S16: STAT5A SH2 domain has two $\beta$-sheet states.** (a) STAT5A trajectories projected on the STAT SH2 conformational landscape. Representative structures from each state are provided (upper: $\beta$D strand in an open state, lower: $\beta$D strand forming contacts in a $\beta$-sheet). (b) Time series of PC2 for each STAT5a trajectory is shown. Each of the six trajectories is indicated in a different colour.

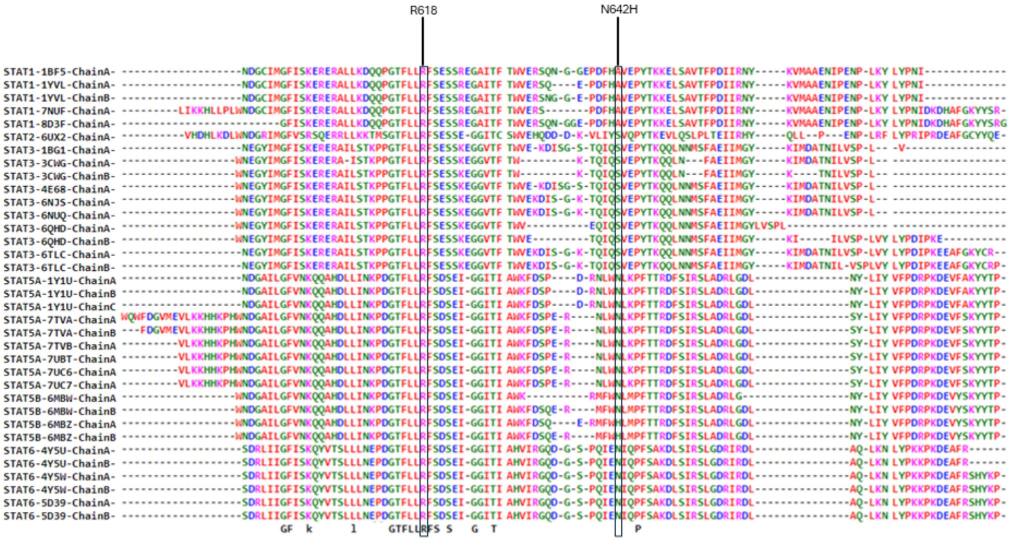


**Figure S17: Sequence alignment of STAT SH2 domains.** Sequence alignment of all mammalian STAT SH2 domains for which a crystal structure is available. The sequence alignment was obtained using MUSTANG [5]. MUSTANG incorporates information from structural alignment based on Cα atoms to determine the sequence alignment [5]. The structures used in the alignment are provided (along with the chain ID from crystal structures containing multiple chains; see Table S2 for the list of crystal structures of mammalian STAT proteins). Residues R618 and N642H (in the residue numbering of STAT5B) are indicated by boxes. Hydrophobic/aromatic residues, acidic, basic, and polar residues are indicated in red, blue, magenta and green, respectively.

| Trajectory | System | # Chains | Length ($\mu$s) |
|---|---|---|---|
| 1 | WT pY-Bound Parallel Dimer | 2 | 2 |
| 2 | WT pY-Bound Parallel Dimer | 2 | 2 |
| 3 | WT pY-Bound Parallel Dimer | 2 | 2 |
| 4 | N642H pY-Bound Parallel Dimer | 2 | 2 |
| 5 | N642H pY-Bound Parallel Dimer | 2 | 2 |
| 6 | N642H pY-Bound Parallel Dimer | 2 | 2 |
| 7 | WT Antiparallel Dimer | 2 | 1.144 |
| 8 | WT Antiparallel Dimer | 2 | 1.144 |
| 9 | WT Antiparallel Dimer | 2 | 1.144 |
| 10 | N642H Antiparallel Dimer | 2 | 2 |
| 11 | N642H Antiparallel Dimer | 2 | 2 |
| 12 | N642H Antiparallel Dimer | 2 | 2 |
| 13 | N642H Antiparallel Dimer | 2 | 1.228 |
| 14 | N642H Antiparallel Dimer | 2 | 1.218 |
| 15 | WT Monomer | 1 | 2 |
| 16 | WT Monomer | 1 | 2 |
| 17 | WT Monomer | 1 | 2 |
| 18 | WT Monomer | 1 | 2 |
| 19 | WT Monomer | 1 | 2 |
| 20 | WT Monomer | 1 | 2 |
| 15 | N642H Monomer | 1 | 2 |
| 16 | N642H Monomer | 1 | 2 |
| 17 | N642H Monomer | 1 | 2 |
| 18 | N642H Monomer | 1 | 2 |
| 19 | N642H Monomer | 1 | 2 |
| 20 | N642H Monomer | 1 | 2 |
| 21 | STAT1 Monomer | 1 | 3 |
| 22 | STAT1 Monomer | 1 | 2.254 |
| 23 | STAT1 Monomer | 1 | 3 |
| 24 | STAT1 Monomer | 1 | 2.256 |
| 25 | STAT1 Monomer | 1 | 2.986 |
| 26 | STAT1 Monomer | 1 | 2.276 |
| 27 | STAT5a Monomer | 1 | 2 |
| 28 | STAT5a Monomer | 1 | 2 |
| 29 | STAT5a Monomer | 1 | 2 |
| 30 | STAT5a Monomer | 1 | 2 |
| 31 | STAT5a Monomer | 1 | 2 |
| 32 | STAT5a Monomer | 1 | 2 |

**Table S1: List of simulations.** The list of all systems in this study is provided along with the length of each simulation used for structural analysis. For the simulations of the wild-type antiparallel dimers, the trajectories that were analyzed included up to 1.144 $\mu$s, which is the time at which the first of these dimers began to dissociate. For the STAT5B$^{N642H}$ antiparallel dimer simulations, three of the simulations did not show dissociation and these were run to 2 $\mu$s. In the other two simulations, the simulations were truncated at the time at which dissociation occurred (1.228 and 1.218 $\mu$s, respectively). Simulations #1-20 are of STAT5B, and simulations #21-32 are of other STAT SH2 domains.

| STAT Protein | PDB ID | Dimerization State |
|---|---|---|
| STAT1 | 1YVL | Antiparallel Dimer |
| STAT1 | 1BF5 | Parallel Dimer |
| STAT1 | 7NUF | Monomer |
| STAT1 | 8D3F | Monomer |
| STAT2 | 6UX2 | Monomer |
| STAT3 | 6NJS | Monomer |
| STAT3 | 6NUQ | Monomer |
| STAT3 | 6TLC | Antiparallel Dimer |
| STAT3 | 6QHD | Parallel Dimer |
| STAT3 | 4E68 | Parallel Dimer |
| STAT3 | 1BG1 | Parallel Dimer |
| STAT3 | 3CWG | Parallel Dimer |
| STAT5A | 1Y1U | Antiparallel Dimer |
| STAT5A | 7UC6 | Monomer |
| STAT5A | 7UC7 | Monomer |
| STAT5A | 7TVA | Antiparallel Dimer |
| STAT5A | 7TVB | Monomer |
| STAT5A | 7UBT | Monomer |
| STAT5B | 6MBW | Antiparallel Dimer |
| STAT5B | 6MBZ | Antiparallel Dimer |
| STAT6 | 4Y5U | Parallel Dimer |
| STAT6 | 4Y5W | Parallel Dimer |
| STAT6 | 5D39 | Parallel Dimer |

**Table S2:** List of crystal structures of mammalian STAT SH2 domains deposited in the PDB as of August, 2024. The STAT SH2 crystal structures in the PDB are 1BF5 [6], 1BG1 [7], 1UUR [8], 1UUS [8], 1Y1U [9](chains A,B,C), 1YVL [10](chains A and B), 3CWG [11](chains A and B), 4E68 [12], 4Y5U [13](chains A and B), 4Y5W [13](chains A and B), 5D39 [13](chains A and B), 6MBW [1](chains A and B), 6MBZ [1](chains A and B), 6NJS [14], 6NUQ [14], 6QHD [15] (chains A and B), 6TLC [16](chains A and B), 7TVA [17] (chains A and B), 7TVB [17], 7UC6 [18], 7UC7 [18], 7UBT [18], 6UX2 [19], 8D3F [20], 7NUF [21]. 1UUR [8] and 1UUS [8] are non-mammalian ancient versions of the STAT protein and are highly dissimilar to the other structures. They were, therefore, excluded from the analysis in this study.

#### References


[1] de Araujo ED, Erdogan F, Neubauer HA, Meneksedag-Erol D, Manaswiyoungkul P, Eram MS, Seo HS, Qadree AK, Israelian J, Orlova A, Suske T, Pham HTT, Boersma A, Tangermann S, Kenner L, Rülicke T, Dong A, Ravichandran M, Brown PJ, Audette GF, Rauscher S, Dhe-Paganon S, Moriggl R, Gunning PT (2019) Structural and functional consequences of the STAT5B N642H driver mutation. Nature Communications 10:2517.

[2] Emenecker RJ, Griffith D, Holehouse AS (2021) Metapredict: a fast, accurate, and easy-to-use predictor of consensus disorder and structure. Biophysical Journal 120(2):4312–4319.

[3] Emenecker RJ, Griffith D, Holehouse AS (2022) Metapredict v2: An update to metapredict, a fast, accurate, and easy-to-use predictor of consensus disorder and structure. bioRxiv p. 10.1101/2022.06.06.494887.

[4] Lim CP, Cao X (2006) Structure, function, and regulation of STAT proteins. Molecular BioSystems 2:536.

[5] Konagurthu AS, Whisstock JC, Stuckey PJ, Lesk AM (2006) MUSTANG: A multiple structural alignment algorithm. Proteins 64(3):559–574.

[6] Chen X, Vinkemeier U, Zhao Y, Jeruzalmi D, Darnell JE, Kuriyan J (1998) Crystal structure of a tyrosine phosphorylated STAT-1 dimer bound to DNA. Cell 93(5):827–839.

[7] Becker S, Groner B, Müller CW (1998) Three-dimensional structure of the STAT3$\beta$ homodimer bound to DNA. Nature 394:145–151.

[8] Soler-Lopez M, Petosa C, Fukuzawa M, Ravelli R, Williams JG, Müller CW (2004) Structure of an activated dictyostelium STAT in its DNA-unbound form. Molecular Cell 13:791–804.

[9] Neculai D, Neculai AM, Verrier S, Straub K, Klumpp K, Pfitzner E, Becker S (2005) Structure of the unphosphorylated STAT5a dimer. Journal of Biological Chemistry 280(49):40782–40787.

[10] Mao X, Ren Z, Parker GN, Sondermann H, Pastorello MA, Wang W, McMurray JS, Demeler B, Darnell JE, Chen X (2005) Structural bases of unphosphorylated STAT1 association and receptor binding. Molecular Cell 17(6):761–771.

[11] Ren Z, Mao X, Mertens C, Krishnaraj R, Qin J, Mandal PK, Romanowski MJ, McMurray JS, Chen X (2008) Crystal structure of unphosphorylated STAT3 core fragment. Biochemical and Biophysical Research Communications 374:1–5.

[12] Nkansah E, Shah R, Collie GW, Parkinson GN, Palmer J, Rahman KM, Bui TT, Drake AF, Husby J, Neidle S, Zinzalla G, Thurston DE, Wilderspin AF (2013) Observation of unphosphorylated STAT3 core protein binding to target dsDNA by PEMSA and X-ray crystallography. FEBS Letters 587:833–839.

[13] Li J, Rodriguez JP, Niu F, Pu M, Wang J, Hung LW, Shao Q, Zhu Y, Ding W, Liu Y, Da Y, Yao Z, Yang J, Zhao Y, Wei GH, Cheng G, Liu ZJ, Ouyang S (2016) Structural basis for DNA recognition by STAT6. Proceedings of the National Academy of Sciences 113:13015–13020.

[14] Bai L, Zhou H, Xu R, Zhao Y, Chinnaswamy K, McEachern D, Chen J, Yang CY, Liu Z, Wang M, Liu L, Jiang H, Wen B, Kumar P, Meagher JL, Sun D, Stuckey JA, Wang S (2019) A potent and selective small-molecule degrader of STAT3 achieves complete tumor regression in vivo. Cancer Cell 36:498–511.

[15] Belo Y, Mielko Z, Nudelman H, Afek A, Ben-David O, Shahar A, Zarivach R, Gordan R, Arbely E (2019) Unexpected implications of STAT3 acetylation revealed by genetic encoding of acetyl-lysine. Biochimica et Biophysica Acta (BBA) - General Subjects 1863(9):1343–1350.

[16] La Sala G, Michiels C, Kükenshöner T, Brandstoetter T, Maurer B, Koide A, Lau K, Pojer F, Koide S, Sexl V, Dumoutier L, Hantschel O (2020) Selective inhibition of STAT3 signaling using monobodies targeting the coiled-coil and n-terminal domains. Nature Communications 11:4115.

[17] Kaneshige A, Bai L, Wang M, McEachern D, Meagher JL, Xu R, Wang Y, Jiang W, Metwally H, Kirchhoff PD, Zhao L, Jiang H, Wang M, Wen B, Sun D, Stuckey JA, Wang S (2023) A selective small-molecule STAT5 protac degrader capable of achieving tumor regression in vivo. Nature Chemical Biology 19:703–711.

[18] Kaneshige A, Bai L, Wang M, McEachern D, Meagher JL, Xu R, Kirchhoff PD, Wen B, Sun D, Stuckey JA, Wang S (2023) Discovery of a potent and selective STAT5 PROTAC degrader

with strong antitumor activity in vivo in acute myeloid leukemia. Journal of Medicinal Chemistry 66:2717–2743.

[19] Wang B, Thurmond S, Zhou K, Sánchez-Aparicio MT, Fang J, Lu J, Gao L, Ren W, Cui Y, Veit EC, Hong H, Evans MJ, O'Leary SE, García-Sastre A, Zhou ZH, Hai R, Song J (2020) Structural basis for STAT2 suppression by flavivirus NS5. Nature Structural & Molecular Biology 27:875–885.

[20] Huang Z, Liu H, Nix J, Xu R, Knoverek CR, Bowman GR, Amarasinghe GK, Sibley LD (2022) The intrinsically disordered protein TgIST from toxoplasma gondii inhibits STAT1 signaling by blocking cofactor recruitment. Nature Communications 13:4047.

[21] Talbot-Cooper C, Pantelejevs T, Shannon JP, Cherry CR, Au MT, Hyvönen M, Hickman HD, Smith GL (2022) Poxviruses and paramyxoviruses use a conserved mechanism of STAT1 antagonism to inhibit interferon signaling. Cell Host & Microbe 30:357–372.
